# The infection cushion: a fungal “weapon” of plant-biomass destruction

**DOI:** 10.1101/2020.06.26.173369

**Authors:** Mathias Choquer, Christine Rascle, Isabelle R Gonçalves, Amélie de Vallée, Cécile Ribot, Elise Loisel, Pavlé Smilevski, Jordan Ferria, Mahamadi Savadogo, Eytham Souibgui, Marie-Josèphe Gagey, Jean-William Dupuy, Jeffrey A Rollins, Riccardo Marcato, Camille Noûs, Christophe Bruel, Nathalie Poussereau

## Abstract

- Grey mold disease affects fruits, vegetables and ornamental plants around the world, causing considerable losses every year. Its causing agent, the necrotrophic fungus *Botrytis cinerea*, produces infection cushions (IC) that are compound appressorial structures dedicated to the penetration of the plant tissues.
- A microarray analysis was performed to identify genes up-regulated in mature IC. The expression data were supported by RT-qPCR analysis performed *in vitro* and *in planta*, proteomic analysis of the IC secretome and mutagenesis of two candidate genes.
- 1,231 up-regulated genes and 79 up-accumulated proteins were identified. They highlight a secretion of ROS, secondary metabolites including phytotoxins, and proteins involved in virulence: proteases, plant cell wall degrading enzymes and necrosis inducers. The role in pathogenesis was confirmed for two up-regulated fasciclin genes. DHN-melanin pathway and chitin deacetylases genes are up-regulated and the conversion of chitin into chitosan was confirmed by differential staining of the IC cell wall. In addition, up-regulation of sugar transport and sugar catabolism encoding genes was found.
- These results support a role for the *B. cinerea* IC in plant penetration and suggest other unexpected roles for this fungal organ, in camouflage, necrotrophy or nutrition of the pathogen.

## Introduction

Many phytopathogenic fungi differentiate specific structures named appressoria that are dedicated to the penetration of the host tissues (Emmet and Parbery, 1975; Deising *et al*, 2000). Appressoria facilitate the breaching of plant cuticles and cell walls through a mechanical and/or chemical action. Early microscopy studies provided clear observations of these structures in different fungal species and distinguished unicellular appressoria (UA) from multicellular appressoria, referred to as infection cushions (IC) (Emmet and Parbery, 1975). Later, UA have attracted much attention and molecular understanding of their development and function has tremendously increased (Ryder and Talbot, 2015). In contrast, IC remain far less understood at the molecular level.

*Botrytis cinerea* is an ascomycetous fungus that causes grey mould disease on more than 1000 plant species (Elad *et al*., 2016). It has been ranked among the ten most severe fungal plant pathogen due to its negative impact on agronomically very important crops and on plant products in post-harvest storage. *B. cinerea* is considered a typical necrotrophic fungus and has become a model to study plant infection (Dean *et al.*, 2012). It is characterized by its ability to produce either UA or IC *in vitro* or *in planta* (Choquer *et al.*, 2007). The *B. cinerea* IC is a compound appressorium that develops between 24 h to 48 h after spore germination.

Histological studies have revealed IC from *B. cinerea* on a wide variety of infected plant hosts and organs: carrot roots (Sharman & Heale, 1977); bean or mung bean hypocotyls (Garcia-Arenal & Sagasta, 1980; Backhouse & Willets, 1987), bean, cucumber or oilseed rape leaves (Akutsu *et al.,* 1981; Van den Heuvel & Waterreus, 1983; Zhang *et al*., 2010), stone fruit or waxflower flowers (Fourie & Holtz, 1994; Dinh *et al*., 2011), lemon or persimmon fruits (Fullerton *et al*., 1999; Rheinländer *et al*., 2013) and onion epidermis (Choquer *et al.*, 2007). Lastly, *B. cinerea* IC also occur *in vitro,* in culture over hard surfaces (Backhouse and Willetts, 1987).

IC have been described in several other Leotiomycetes fungi: *Sclerotinia sclerotiorum* (Tariq & Jeffries, 1984), *Sclerotinia minor* (Lumsden & Wergin, 1980)*, Sclerotinia trifoliorum* (Prior & Owen, 1964)*, Dumontinia tuberosa* (Pepin R, 1980), and *Stromatinia cepivora* (Stewart *et al*., 1989). In parallel, IC were described in the distant ascomycota *Tapesia yallundae* (Daniels *et al*., 1991), *Fusarium graminearum* (Boenisch & Schäfer, 2011), and in the basidiomycota *Athelia rolfsii* (Smith *et al*., 1986), *Rhizoctonia solani* (Demirci & Döken, 1998) and *Rhizoctonia tuliparum* (Gladders & Coley-Smith, 1977). More recently, IC-related structures have also been observed in several saprophytic fungi highlighting the role of these structures in biomass degradation (Demoor *et al*., 2019).

The characterization of several avirulent mutants of *B. cinerea* revealed the importance of IC in the infectious process of this necrotrophic fungus (De Vallée *et al*., 2019) and hypovirulent strains of *B. cinerea* infected by mycoviruses are deficient in IC formation (Zhang *et al*., 2010; Hao *et al*., 2018). Although it has long been accepted that IC mediate penetration, their differentiation as well as their functions are still poorly understood. By using a transcriptomic approach in *B. cinerea*, the aim of this study was to provide new molecular information that highlights biological processes specifically at work in the mature IC. These structures were differentiated on a synthetic membrane mimicking the plant surface and gene expression was compared between a mycelium enriched in IC and a control mycelium produced under agitated culture conditions.

The transcriptomic results, supported by a secretome analysis, are consistent with IC being structures dedicated to the secretion of fungal effectors important for plant penetration and colonization: phytotoxins, ROS, hydrolytic enzymes and plant necrosis inducers. Moreover, the data reveal a deep remodeling of the IC cell wall suggesting the importance of melanin and chitosan in the function of IC. The hypothesis of a role in the nutrition of the parasite is also proposed for the IC.

## Materials and Methods

### Fungal strains and growth conditions

*Botrytis cinerea* B05.10 conidia were collected in PDB medium (Difco) diluted ¼ (PDB¼), after ten days of culture on malt sporulation medium, at 21°C under near-UV light. All subsequent cultures were done in the dark at 21°C. For infection cushions formation, 2.10^5^ conidia were spread onto cellophane sheets placed on PDB¼ medium supplemented with agar (25 g.l^−1^), then incubated for 44 h. Simultaneously, same quantity of conidia was inoculated in 250 ml-flasks containing 50 ml PDB¼ medium and incubated for 44 h under 110 rpm agitation. Flasks were previously siliconized with Sigmacote (Sigma) to prevent mycelium adhesion and IC formation on the glassware during culture. For radial growth measurements, wild-type and mutant strains were grown on minimal medium (NaNO_3_ 2 g.l^−1^, Glucose 20 g.l^−1^, KH_2_PO_4_ 0.2 g.l^−1^, MgSO_4_, 7H_2_O 0.1 g.l^−1^, KCl 0.1 g.l^−1^, FeSO_4_, 7H_2_O 4 mg.l^−1^). For secretome analysis, 2.10_5_ conidia were spread onto cellophane membrane overlaying solid PDB¼. After 6 h incubation (21°C), membranes were transferred on 2 ml liquid PDB¼ for 24h (Control mycelium condition) or 48 h (IC condition) at 21°C. Liquid medium was then collected for proteomic analysis.

### Microarray expression analysis

To study the transcriptome of *B. cinerea*, NimbleGen 4-plex arrays containing 4 × 72,000 arrays per slide were used (Roche, Mannheim, Germany). Construction of this chip was initially based on combining two previous genome annotations (Amselem *et al*., 2011), the one of the T4 strain by URGI (BofuT4 gene references; www.urgi.versailles.inra.fr) and the one of the B05.10 strain by the Broad Institute (BC1G gene references; www.broadinstitute.org). Thus, 62,478 60-mer oligonucleotides were designed as specific probes covering 20,885 predicted gene models and non-mapping Expressed Sequence Tags (EST) (3 oligonucleotides per gene or EST) and 9,559 random probes were designed as negative controls.

The structural annotation used for this study was published by van Kan *et al.* (2017), displaying 11,718 predicted genes, associated to 13,749 predicted proteins, unlike the two previous annotations showing more than 16,000 predicted genes (BofuT4 and BC1G). This annotation is considered better on the basis of RNA-seq data and is available at EnsemblFungi under the reference *Botrytis cinerea* B05.10 (ASM83294v1; Bcin gene references; http://fungi.ensembl.org/Botrytis_cinerea/). 11,718 Bcin genes are predicted but not all of them were analyzed by the *B. cinerea* NimbleGen 4-plex arrays. In order to associate each BofuT4 and BC1G gene from the chip to only one Bcin gene and reciprocally, we used the correspondence established in the *B. cinerea* Portal (Simon & Viaud, 2018; http://botbioger.versailles.inra.fr/botportal/). In a hundred of cases, BofuT4/BC1G genes showing correspondence with multiple Bcin genes were checked by gene synteny. After this manual curation, we identified 11,630 BofuT4/ BC1G genes out of 15,750 showing correspondence with only one Bcin gene. When a Bcin gene showed correspondence with multiple BofuT4/ BC1G genes, we selected the BofuT4 or BC1G gene giving the highest normalized intensities. After these two manual curations, we found that 11,134 Bcin genes were analyzed by the chip which represent 95% of the published Bcin genes (11,718; Table S1). The EST and the small coding sequences (< 100 amino acids) lacking EST support were excluded from our analysis.

Total RNA was extracted from 4 mg of ground lyophilized material using the RNeasy Midi kit (Qiagen). A DNase treatment (Ambion) was performed to remove traces of genomic DNA. RNA profiles were assessed using the Bioanalyzer RNA 6000 Nano kit (Agilent). Ten micrograms of total RNA were converted into cDNA using the SuperScript II cDNA Conversion Kit (Invitrogen). Double stranded cDNAs were then labelled with Cy3-nonamers using NimbleGen One-Color DNA Labeling Kit before hybridization on the NimbleGen 4-plex arrays. Microarrays were then scanned with an Agilent scanner at 532 nm (Cy3 absorption peak) optimized for Nimblegen 4-plex arrays. All steps were performed following the procedures established by NimbleGen. The entire microarray data set described in this article is available at the Gene Expression Omnibus (GEO) database under accession number GSE141822.

Data processing, quality controls, differential expression analysis and clustering were performed using ANAIS methods (Simon and Biot, 2010). Hybridization signals of all probes, comprising three and four independent replicates for mycelium condition and infection cushion condition, respectively, were subjected to RMA-background correction, quantile normalization, and gene summarization. Thresholds of gene expression were determined by referring the hybridization signals to those of 9,559 random probes, calculated for each array using the R software (R Core Team, 2013). Genes were considered expressed when their normalized intensity was higher than the 99th percentile of random probes hybridization signals in at least one biological replicate. These genes were kept for differential expression analysis. Differentially expressed genes, between the infection cushion condition and the mycelium condition, were identified using a one-way ANOVA test. To deal with multiple testings, the ANOVA p-values were submitted to a False Discovery Rate correction (FDR). Transcripts with a corrected p-value <0.05 and for which a fold change ≤ −2 or ≥ 2 was observed between the two conditions were considered to display significant differential expression. Clusters analyses of gene normalized intensities were performed to highlight differentially expressed genes.

### Enrichment analysis and genes categorization

Further analyses were performed to highlight biological processes potentially enriched in the selected lists of up-regulated or down-regulated Bcin genes in infection cushion. Enrichment in Gene Ontology (GO) Biological Process (BP) terms was assessed on the *Botrytis cinerea* B05.10 (ASM83294v1) species using the Fungifun website (Priebe *et al*., 2015; https://elbe.hki-jena.de/fungifun/). The background dataset of genes used as reference was the 11134 Bcin genes which were associated to the chip (Table S1). Presence of a putative signal peptide was predicted using the SignalP 5.0 Server (Almagro Armenteros *et al.*, 2019). The CAZy database (www.cazy.org) was used for the manual curation of plant cell wall degrading enzymes and the alpha-1,2-mannosidases. The InterPro database (www.ebi.ac.uk/interpro/; Mitchell *et al.*, 2019) was used for the manual curation of cytochrome P450s, taurine catabolism dioxygenases, cutinases, sugar transporters, sedolisins, serine carboxypeptidases and metallopeptidases.

### Expression profiling by quantitative PCR analysis

Experiments were performed as described by Rascle *et al*. (2018). RT-qPCR experiments were performed using ABI-7900 Applied Biosystems (Applied Biosystems). Amplification reactions were carried out using SYBR Green PCR Master Mix (Applied Biosystems). Relative quantification was based on the 2(−ΔΔC(T)) method (Livak & Schmittgen, 2001) using the *BcactA* (Bcin16g02020), *Bcef1α* gene (Bcin09g05760), and *Bcpda1* gene (Bcin07g01890) as normalization internal controls. At least three independent biological replicates were analyzed. Primers used for RT-qPCR are shown in Table S5.

### ROS and melanin production

For visualization of melanin, a stock solution of the DHN melanogenesis inhibitor tricyclazole (Sigma) was prepared in acetone (10 mg ml^−1^). Three-day-old 2-mm mycelial plugs were deposited in 6-well plates. Droplets (50 μl) of PDB¼ (supplemented or not with 50 μg ml^−1^ tricyclazole) medium were added on the plugs. After 48 h incubation (21°C), IC were observed by reverse microscopy. For visualization of ROS produced during IC formation, 100μl of medium were added to 1 ml of DAB (Sigma) solution (0.05% in 100 mM citric acid buffer pH 3.7) and incubated 20 h in darkness with gentle agitation. Controls were done by adding 1 μl Horse Radish Peroxydase (HRP - Thermoscientific) to 1 ml DAB in presence of different quantities of H_2_O_2._

### Proteomic analysis

The steps of sample preparation and protein digestion were performed as previously described (Dieryckx *et al.*, 2015) and online nanoLC-MS/MS analyses were performed using an Ultimate 3000 RSLC Nano-UPHLC system (Thermo Scientific) coupled to a nanospray Q Exactive hybrid quadrupole-Orbitrap mass spectrometer (Thermo Scientific). The parameters of the LC-MS method used were as previously described (Pineda *et al.*, 2018). Protein identification and Label-Free Quantification (LFQ) were done in Proteome Discoverer 2.3. MS Amanda 2.0, Sequest HT and Mascot 2.4 algorithms were used for protein identification in batch mode by searching against the Ensembl *Botrytis cinerea* B05.10 database (ASM83294v1, 13 749 entries, release 98.3). Two missed enzyme cleavages were allowed. Mass tolerances in MS and MS/MS were set to 10 ppm and 0.02 Da. Oxidation (M), acetylation (K) and deamidation (N, Q) were searched as dynamic modifications and carbamidomethylation (C) as static modification. Peptide validation was performed using Percolator algorithm (Käll *et al.*, 2007) and only “high confidence” peptides were retained, corresponding to a 1% false discovery rate at peptide level. Minora feature detector node (LFQ) was used along with the feature mapper and precursor ions quantifier. The normalization parameters were selected as follows: (1) Unique peptides, (2) Precursor abundance based on intensity, (3) Normalization mode: total peptide amount, (4) Protein abundance calculation: summed abundances, (5) Protein ratio calculation: pairwise ratio based and (6) Hypothesis test: t-test (background based). Quantitative data were considered for master proteins, quantified by a minimum of 2 unique peptides, a fold change ≥3 and a statistical p-value lower than 0.05. The mass spectrometry proteomics data have been deposited to the ProteomeXchange Consortium (http://proteomecentral.proteomexchange.org) via the PRIDE partner repository (Perez-Riverol *et al*., 2019) with the dataset identifier PXD016885.

### Construction of deletion mutants in *Botrytis cinerea*

Δ*Bcflp1* and Δ*Bcflp2* deletion mutants were constructed using a gene replacement strategy (Fig.S5). The replacement cassettes were generated by combining double-joint PCR (Yu *et al*., 2004) and split-marker approach (Catlett *et al.*, 2003). All primers are listed in Table S4. The three gene replacement cassettes were verified by sequencing. *B. cinerea* transformation was carried out using protoplasts as previously described by Lalève *et al.* (2014), except that the protoplasts were transformed with 1μg of each split-marker cassette DNA and plated on medium containing 200 g.l^−1^ saccharose and 2 g.l^−1^ NaNO_3_ supplemented with 70 μg.ml^−1^ hygromycin (Invivogen, France) for single Δ*Bcflp1* or Δ*Bcflp2* mutants or 80 μg.ml^−1^ nourseothricin (Werner BioAgents, Germany) for double replacement mutants. Diagnostic PCR was performed to detect homologous recombination in the selected resistant transformants. Homokaryotic transformants were obtained after several rounds of single-spore isolation. Southern blot analyses were performed to ensure single insertions. Genomic DNA digested with *Eco*RI was hybridized with the 3’-flanking region of *Bcflp1* (amplified with Probe1For and Probe1Rev primers) or the 5’-flanking region of *Bcflp2* (amplified with Probe2For and Probe2Rev primers) using the PCR DIG Probe Synthesis Kit and the DIG Luminescent Detection Kit (Roche, Germany) following manufacturer’s instructions.

### Pathogenicity tests

Infection assays were performed with one-week-old French bean (*Phaseolus vulgaris* var *Saxa*) leaves using 4-mm 72-h-old mycelial plugs from *B. cinerea* WT, Δ*Bcflp1*, Δ*Bcflp2* and Δ*Bcflp1*::Δ*Bcflp2* mutant strains grown on sporulation medium. Infected plants were incubated at 21°C under 100% relative humidity and dark (10 h)-daylight (14 h) conditions. Necrosis zone diameter was measured daily. All tests were assessed in three independent experiments, with at least eight plants and thirty points of infection for each strain.

## Results

### Microarray study of IC

To gain more information on the role of IC in the biology of *B. cinerea*, we identified genes expressed in these structures. As IC develop on solid surfaces, agar plates overlaid with cellophane sheets were used for their production while a control vegetative mycelium was prepared in agitated liquid cultures. Conidia were used to inoculate both solid and liquid potato dextrose media. To obtain as much IC as possible, the samples were collected at 44 hpi. At that time, IC were fully differentiated and hyperbranched (Fig. 1), and they covered about 40% of the PDA plates surfaces (Fig. S1a) while the liquid cultures produced only mycelium (Fig. S1b). Both biological materials were used to extract total RNA and to prepare cDNAs that were hybridized to microarrays carrying probes of 11,134 *B. cinerea* genes (Table S1). Data processing, quality controls and differential expression analysis of the microarrays data led to the listing of 1,231 up-regulated genes and 1,422 down-regulated Bcin genes (fold change ≤ −2 or ≥ 2; FDR < 0.05; Table S1) in the IC-enriched sample (hereafter referred to as IC) when compared to vegetative mycelium (13% and 15% of the 9,410 expressed genes, respectively). The reliability of this result was tested by RT-qPCR analysis. For this, new IC and new control vegetative mycelium were produced in order to extract new RNAs and to run the qPCR reactions on 39 genes selected among the up-, down- and non-regulated genes identified by the microarray study (the selection covered genes with different fold-changes). The results confirmed the microarrays data in 85% of the cases (33 genes) and showed no opposite expression profile between the microarray and RT-qPCR results for all the up- and down-regulated genes (Table S2). Altogether, these results granted the microarray data a good level of confidence.

**Figure 1:**
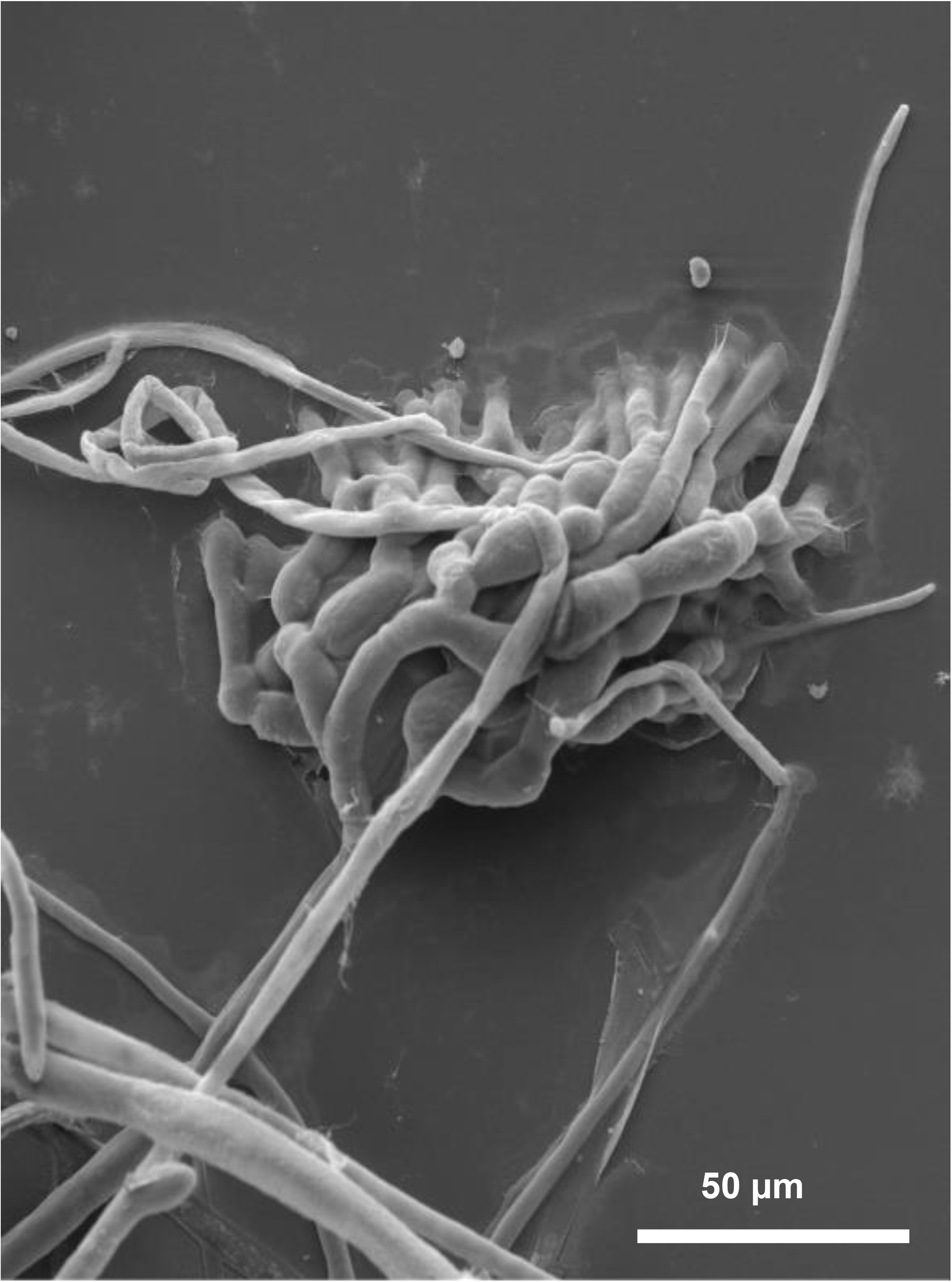
Scanning Electron Microscopy of *Botrytis cinerea* infection cushion. Mature infection cushion, produced on glass surface, developing multiple and successive ramifications of vegetative hyphae (2 dpi).

### Functional enrichment analysis of differentially expressed genes in IC

Enrichment analyses was performed on the up- and down-regulated genes in IC. By using the GO biological process classification (Gene Ontology; Fig. S2), a significant enrichment was revealed for carbohydrate metabolic process (58 genes), oxidation-reduction process (136 genes), metabolism process (70 genes), transmembrane transport (76 genes) and proteolysis (24 genes). For down-regulated genes, rRNA processing (15 genes), ribosome biogenesis (9 genes) and tRNA splicing (5 genes) showed enrichment. In order to extract biological meaning from the GO analysis, we used existing databases and published data to manually sort the transcriptional data into 21 functional subcategories (Table S1). Fisher’s exact tests showed the enrichment of 17 of these 21 subcategories, relating to plant degradation, fungal cell wall remodeling, production of secondary metabolites, phytotoxins or ROS, and potentially nutrition and secretion (Table 1).

**Table 1:**
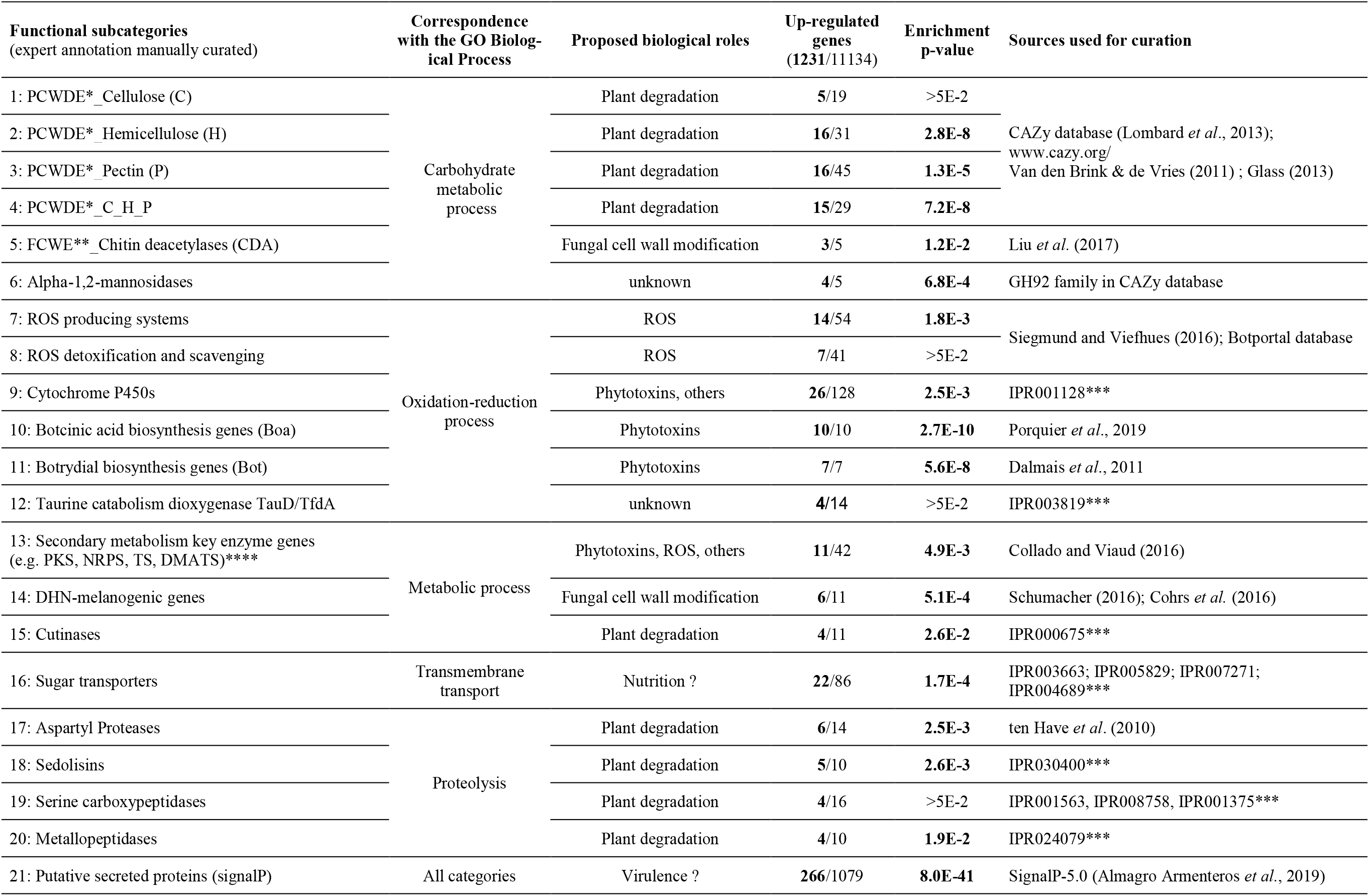
Functional enrichment analysis of up-regulated genes in the infection cushion of *Botrytis cinerea*. Functional subcategories were determined according to a GO biological process analysis followed by a manual curation using existing databases and published data. Categories with significant enrichment were identified using Fisher’s exact test with a p-value cutoff at 0.05; Number of differentially regulated genes (in bold) and p-values are indicated. *PCWDE, Plant Cell Wall Degrading Enzymes; **FCWE, Fungus Cell Wall Enzymes; *** Bcin proteins displaying an “IPR” InterPro domain (www.ebi.ac.uk/interpro/; Mitchell *et al.*, 2019) were searched on the Botportal database (http://botbioger.versailles.inra.fr/botportal/); **** PKS, Polyketide synthases; NRPS, Non-Ribosomal Peptide Synthetases; TS, Terpen cyclases; DMATS, Dimethylallyl tryptophan synthases.

### Protein effectors: enzymes degrading the plant barriers and tissues

The carbohydrate-active enzymes database (CAZy, Lombard *et al.*, 2013) and the classifications of Plant Cell Wall Degrading Enzymes (PCWDE; Van den Brink & de Vries, 2011; Glass *et al.*, 2013) were used to sub-classify 126 predicted PCWDE-encoding genes in *B. cinerea* according to their putative specificity for cellulose (C), hemicellulose (H), pectin (P) or overlapping specificity (H, P or C) (Table S1). As shown in Table 1, genes coding for hemicellulases, pectinases, and PCWDE of overlapping specificity were significantly enriched among the up-regulated genes in IC, while the genes coding for cellulases were not. Genes coding for cutinases and for proteases were also found significantly enriched among the up-regulated genes in IC (Table 1). In particular, the aspartyl proteases (Ten Have *et al*., 2010), sedolisins and metallopeptidases sub-categories were enriched. Altogether, these results suggest a role for IC in the degradation of the plant barriers and tissues through the massive production and secretion of hydrolytic enzymes.

### Small molecule effectors: phytotoxins and reactive oxygen species (ROS)

Detailed examination of the predicted 42 genes that code for Secondary Metabolism (SM) key enzymes in *B. cinerea* B05.10 strain (Table S1; Collado & Viaud, 2016) showed that 11 were up-regulated in IC, and this made this sub-category enriched (Table 1; Fig. S3). Noticeably, two SM gene clusters, responsible for the productions of botcinic acid and botrydial phytotoxins (Dalmais *et al.*, 2011; Porquier *et al.*, 2019), were both significantly enriched among the up-regulated genes in IC (subcategories “10” and “11” in Table S1; Table 1; Fig. 2a). In addition, the *Bcreg1* gene (Bcin03g07420) encoding the transcriptional regulator required for the synthesis of these toxins (Michielse *et al.*, 2011) was also up-regulated (Table S1). At last, we observed that five remaining up-regulated SM key enzyme-encoding genes are neighbored by genes whose expression was also up-regulated in IC, and whose description is compatible with SM typical genes (Fig. S4). This suggests the existence of four additional putative SM clusters transcriptionally induced in *B. cinerea* IC. These gene clusters include *Bcdtc3*, *Bcpks7*, *Bcpks8* or *Bcpks4*/*Bcnrps8* genes and the metabolites they would produce are still unknown.

**Figure 2:**
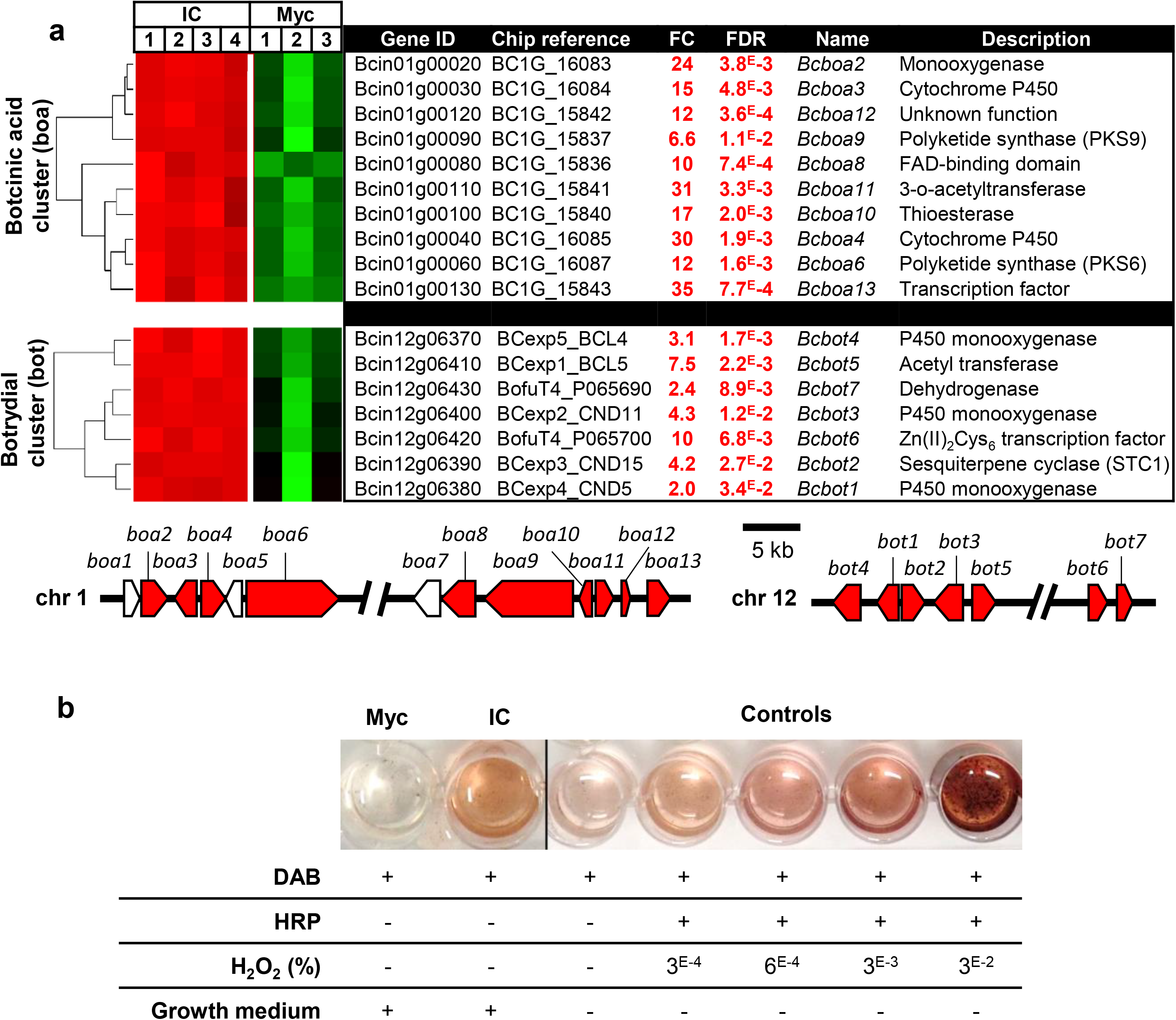
Up-regulation of phytotoxins and ROS production in the infection cushion of *Botrytis cinerea*. (a) Hierarchical clustering of the expression of the Botcinic acid genes (*Bcboa)* and botrydial genes (*Bcbot)* in infection cushion. (IC, 4 biological replicates) and control mycelium (Myc, 3 biological replicates). The normalized expression intensities are clustered and represented by color-coded squares; Shades of green and red depict down- and up-regulation in IC, respectively (fold change (FC) ≤ −2 or ≥ 2 and FDR<0.05). (b) Botcinic acid (boa) and botrydial (bot) structures (left; Pinedo *et al*., 2016) and biosynthesis gene clusters (right; Porquier *et al*., 2019 and Dalmais *et al*., 2011). Genes up-regulated in IC are colored in red. Genes with no chip reference and whose expression could hence not be measured are colored in white. (c) Visualization of ROS produced by IC. 100 μl of medium were added to 1ml of DAB and incubated 20h. Controls were done by adding 1μl Horse Radish Peroxydase (HRP) to 1ml DAB in presence of different quantities of H_2_O_2_. The growth medium of 48 h cellophane-overlaid liquid cultures (IC) or agitated liquid cultures (Myc) was mixed to a DAB solution. The oxidation of DAB by H_2_O_2_ in the presence of peroxidases is revealed by a brown coloration.

Detailed examination of the predicted 54 genes that code for ROS producing systems in *B. cinerea* (Siegmund & Viefhues, 2016) showed that 14 were up-regulated in IC, making this gene family an enriched sub-category (Table 1; Table S1). These genes encode the catalytic subunit BcNoxB of the NADPH oxidase and its regulator BcNoxR (Segmüller *et al.*, 2008), the glucose oxidase BcGod1 (Rolke *et al.*, 2004), the galactose oxidase BcGox1, the quinone oxidoreductase BcNqo1 (An *et al.*, 2016), as well as 3 laccases (BcLcc6, 8, 13) and 6 glucose-methanol-choline (GMC) oxidoreductases. As ROS play a role in the early phases of plant infection by *B. cinerea* (Govrin and Levine, 2000), up-regulation of these genes suggested that IC could secrete more ROS than vegetative mycelium. To test this hypothesis, *B. cinerea* was cultured for 48 h on liquid PDB medium overlaid with cellophane sheets and the collected culture medium was exposed to 3,3’ diaminobenzidine (DAB) as previously described (Viefhues *et al*., 2014). In comparison with the vegetative mycelium, a stronger oxidation of DAB was observed with the IC-culture medium (Fig. 2b), indicating the presence of likely higher H_2_O_2_ concentration and secretion of peroxidase activity.

### Melanization and chitin deacetylation remodel the IC cell wall

In *B. cinerea*, dihydroxynaphthalene-(DHN)-melanins are produced and their biosynthesis relies on a bipartite pathway operating in conidia or in sclerotia (Schumacher, 2016). In conidia, the genes coding for the polyketide synthase BcPks13 and for the hydrolase BcYgh1 are induced by the light transcription factor BcLft2. In sclerotia, the polyketide synthase BcPks12 encoding gene is induced by the transcription factor SMR1 and repressed by BcLtf2 (Cohrs *et al.*, 2016). In IC, *Bcpks13*, *Bcygh1*, *Bcltf2* and the other downstream genes of the pathway were up-regulated while *Bcpks12* and *Bcsmr1* were down-regulated (Fig. 3a-c). This would indicate that IC produce DHN-melanins by using the biosynthetic pathway at play in conidia. Schumacher (2016) proposed laccases as potential candidates catalyzing the polymerization of DHN-melanins monomers in *B. cinerea*. As *BcLcc6* and *BcLcc8* laccases genes are up-regulated in the IC, they could eventually play this role (Sapmak *et al.*, 2015). In addition, thick dark cell walls were observed in IC (Fig. 3d, top), suggesting that DHN-melanins are deposited at the contact zones between the hyperbranched IC lobes. In the presence of tricyclazole, an inhibitor of the THN reductases BcBrn1 and BcBrn2 (Fig. 3b), hyperbranched IC lobes were still formed but their dark cell walls were not visible anymore. Orange cell walls were observed instead (Fig. 3d, bottom), likely due to accumulation of T4HN and T3HN and to their autooxidation products flaviolin and 2-hydroxyjuglone (Schumacher, 2016). These results confirm the hyper-melanization of the IC cell wall and suggest that melanization and hyperbranching are independent processes.

**Figure 3:**
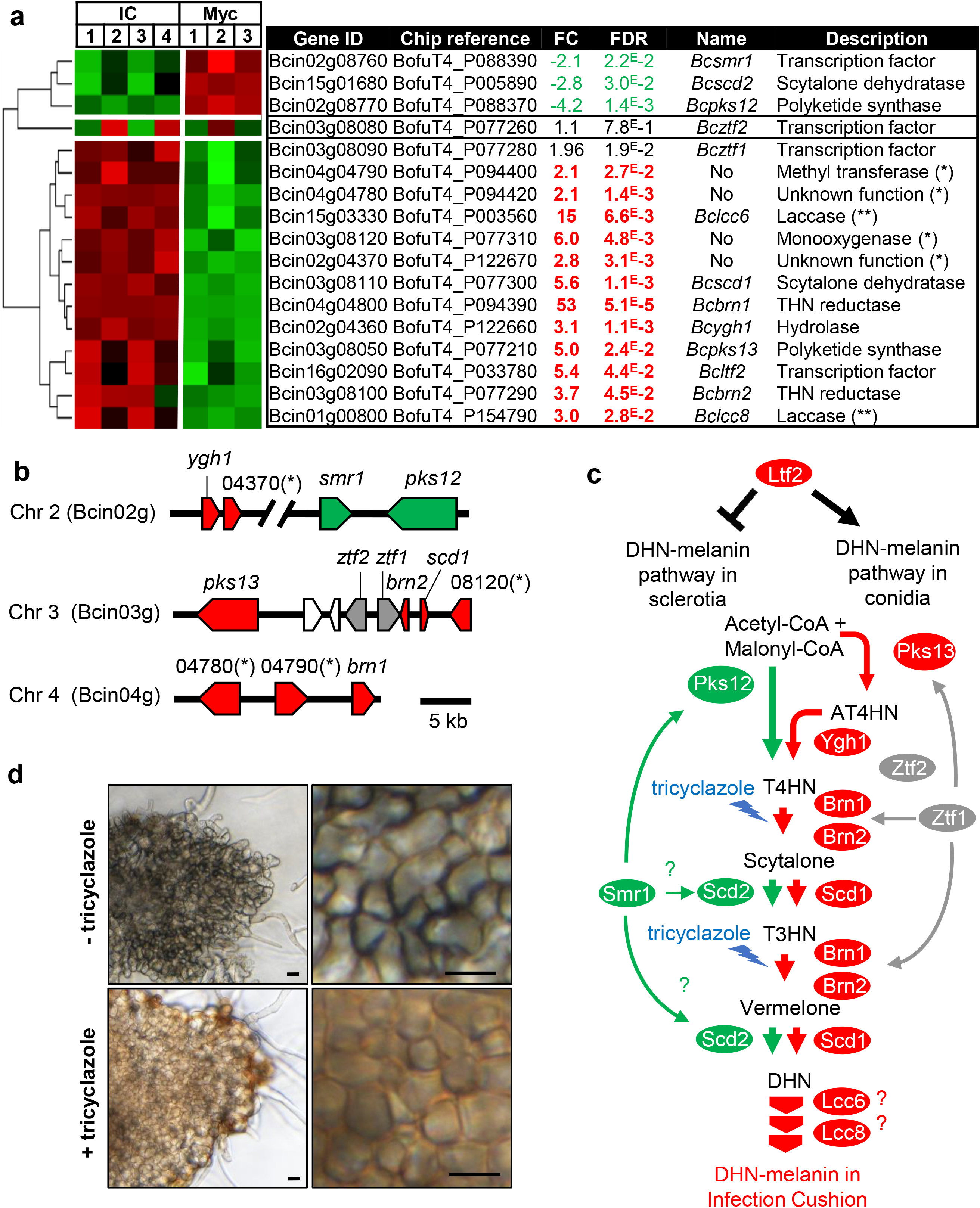
Regulation of the DHN melanogenesis bipartite pathway in the infection cushion of *Botrytis cinerea.* This figure is adapted from Schumacher *et al*. (2016) and Cohrs *et al*. (2016). (a) Hierarchical clustering of the expression of DHN-melanogenic genes in IC (IC, 4 replicates) and control mycelium (Myc, 3 replicates) as presented in figure 2. Up-regulated genes that co-localize with the DHN-melanogenic genes are added for information (*), but their role in melanin biosynthesis remains to be established. Similarly, 2 laccases up-regulated genes (*Bclcc6* and *Bclcc8*) are listed (**), but their role in melanin biosynthesis remains to be established. (b) DHN-melanins biosynthesis putative gene clusters. (c) DHN-melanin metabolic pathway and targets of tricyclazole (lightnings). Genes up- or down-regulated in IC and corresponding proteins are colored in red and green, respectively. (d) Light microscopy images of hyphae and IC of *B. cinerea* produced on plastic surface in the absence (top) and presence (bottom) of tricyclazole. Pigmented IC versus hyaline hyphae (left) and dark versus orange-brown thick cell walls inside IC (right) are shown. Bars represent 10 μm.

Chitin deacetylases (CDA) are enzymes that produce chitosan from chitin (Fig. 4a). Based on conserved protein motifs identified in CDAs (Liu *et al.*, 2017), the genome of *B. cinerea* putatively encodes five CDA (data not shown; Table S1). All these genes were expressed in IC and 3 of them were up-regulated in IC when compared to the control mycelium (Fig. 4b). *In planta* RT-qPCR analysis showed up-regulation of these 5 genes at the early phase of bean leaves infections (Fig. 4c). Whether the up-regulation of the CDA genes impacted the IC cell wall was addressed by using confocal microscopy and differential staining of chitin and chitosan. This allowed the specific visualization of chitosan in IC, hooks and their generating hyphae, but not in the vegetative mycelium (Fig. 4d).

**Figure 4:**
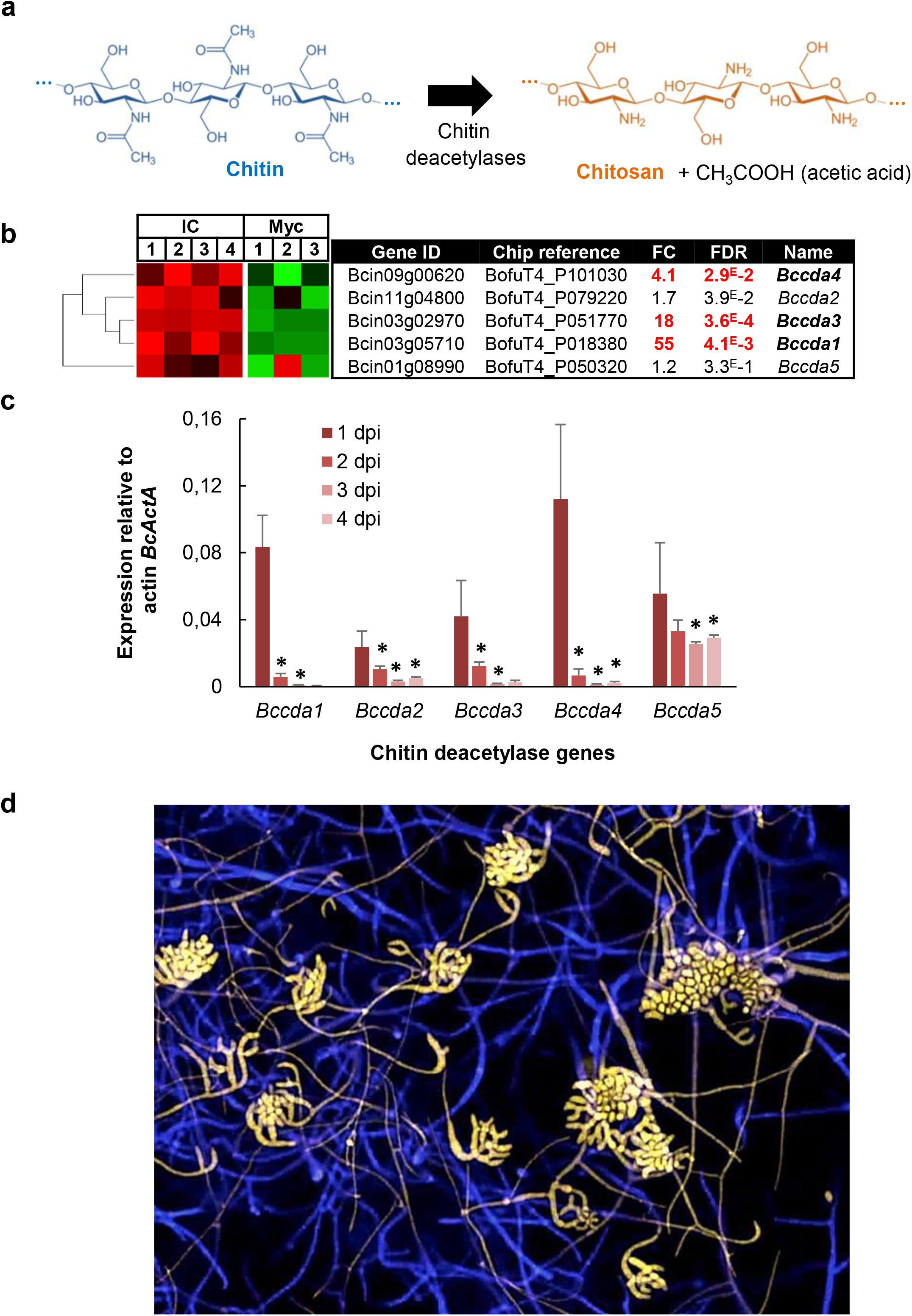
Cell wall chitin deacetylation in the infection cushion of *Botrytis cinerea*. (a) Representation of the transformation of chitin into chitosan by chitin deacetylases. (b) Hierarchical clustering of the expression of 5 putative *cda* genes in IC (IC) and control mycelium (Myc), as presented in figure 2. (c) Expression of *Bccda* genes during a kinetics of bean leaves infection by *B. cinerea* (dpi; days post inoculation). Expression levels were calculated following the 2^−ΔΔCT^ method using constitutively expressed actin gene *BcactA* (Bcin16g02020) as a reference. The use of two other housekeeping genes, elongation factor *Bcef1α* (Bcin09g05760) and pyruvate dehydrogenase *Bcpda1* (Bcin07g01890) gave similar results (data not shown). Three independent biological replicates were assessed for each experiment. Standard errors are displayed, and asterisks indicate a significant difference in gene expression compared with the previous time point (Student’s *t* test, * p-value <0.05). (d) Confocal microscopy of mycelium and IC produced onto a plastic surface and double-stained with Calcofluor targeting mainly chitin (blue) and Eosin Y targeting mainly chitosan (yellow).

### Upregulation of sugar uptake and catabolism in IC

The “Sugar transporters” subcategory was explored by mining the BotPortal database (http://botbioger.versailles.inra.fr/botportal) using the four InterPro domains IPR003663, IPR005829, IPR007271, and IPR004689. This led to the listing of 86 predicted sugar transporters in the genome of *B. cinerea* (Table S1), among which 22 were up-regulated in IC (Table 1). This up-regulation suggests a possible activation of the downstream sugar catabolic pathways in IC. Noticeably, two genes of the glycolysis pathway are up-regulated in IC, coding for the fructose-bisphosphate aldolase (Bcin07g03760; Table S1) and the glucose-6-phosphate isomerase *Bcpgi* (Bcin15g04970; Table S1). Besides, in the gluconeogenesis pathway, the *BcPck1* gene (Bcin16g00630; Table S1) encoding the phosphoenolpyruvate carboxykinase gene is also up-regulated in IC, and Liu *et al*. (2018) showed that this key gene is crucial for *B. cinerea* virulence and for the formation of IC in the absence of glucose. In addition to the enriched metabolic pathways that relate to the degradation of polysaccharides (pectin, glycans), these results suggest an activated sugar catabolism in IC.

### IC secretome analysis validates the transcriptome analysis and highlights the secretion of necrosis-inducing factors

Of the 1,231 up-regulated genes in IC, 266 were predicted to code for putative secreted proteins with a N-terminal signal peptide (Table S1), and this made this sub-category enriched in IC (Table 1). To support this data, the secretomes of IC and control mycelium were compared. *B. cinerea* was grown on cellophane sheets overlaying liquid PDB medium and the culture media were collected at 24 h (control mycelium) or at 48 h when IC covered ~40% of the cellophane surface. The proteins were extracted and subjected to a comparative proteomics analysis. 79 proteins identified in 3 biological replicates with a minimum of 2 unique peptides were listed as up-accumulated (Fold change ≥ 3, FDR < 0.05) in IC (Table S3). These proteins are essentially involved in carbohydrate metabolic process (29 proteins), proteolysis (16 proteins) and oxidation-reduction process (12 proteins), 3 of the 5 functional categories resulting from the GO enrichment analysis of the genes up-regulated in IC (Fig. S2). Besides, comparison of the proteomics and microarray data (Table S3) showed that 40 of the up-accumulated proteins (51%) are encoded by genes up-regulated in IC. The proteomics data therefore support the transcriptomic data and validate the proposed role of IC in enzymes secretion. In addition, 6 known necrosis inducers were up-accumulated in the secretome of IC: BcGs1 (Zhang *et al*., 2015), BcXyn11A (Brito *et al*., 2006), BcXyg1 (Zhu *et al.*, 2017), BcSpl1 (Frias *et al*., 2013), BcNep2 (Schouten *et al.*, 2008) and BcIeb1 (Frias *et al*., 2016). This suggests that IC could actively participate in the process of plant cell necrosis.

### Relevance of the microarray data to plant infection

Fourteen up-regulated genes in IC whose predicted function relates to pathogenesis were selected for a time course RT-qPCR analysis *in planta* (bean leaves infected with conidia of *B. cinerea*; Fig. 5a and 5b). All the selected genes were expressed *in planta* suggesting that the microarray data are relevant to plant infection. Twelve of the studied genes were upregulated at days 2, 3 and/or 4 of infection, when IC were visible at the leaf surface. When only hooks could be observed (day 1), most of these genes (11) were not expressed. Furthermore, expression peaked at day 2 for 7 genes, when IC were visible, but plant tissue maceration was not. These results would indicate that these proteins secreted early by IC could participate to the plant cell death.

**Figure 5:**
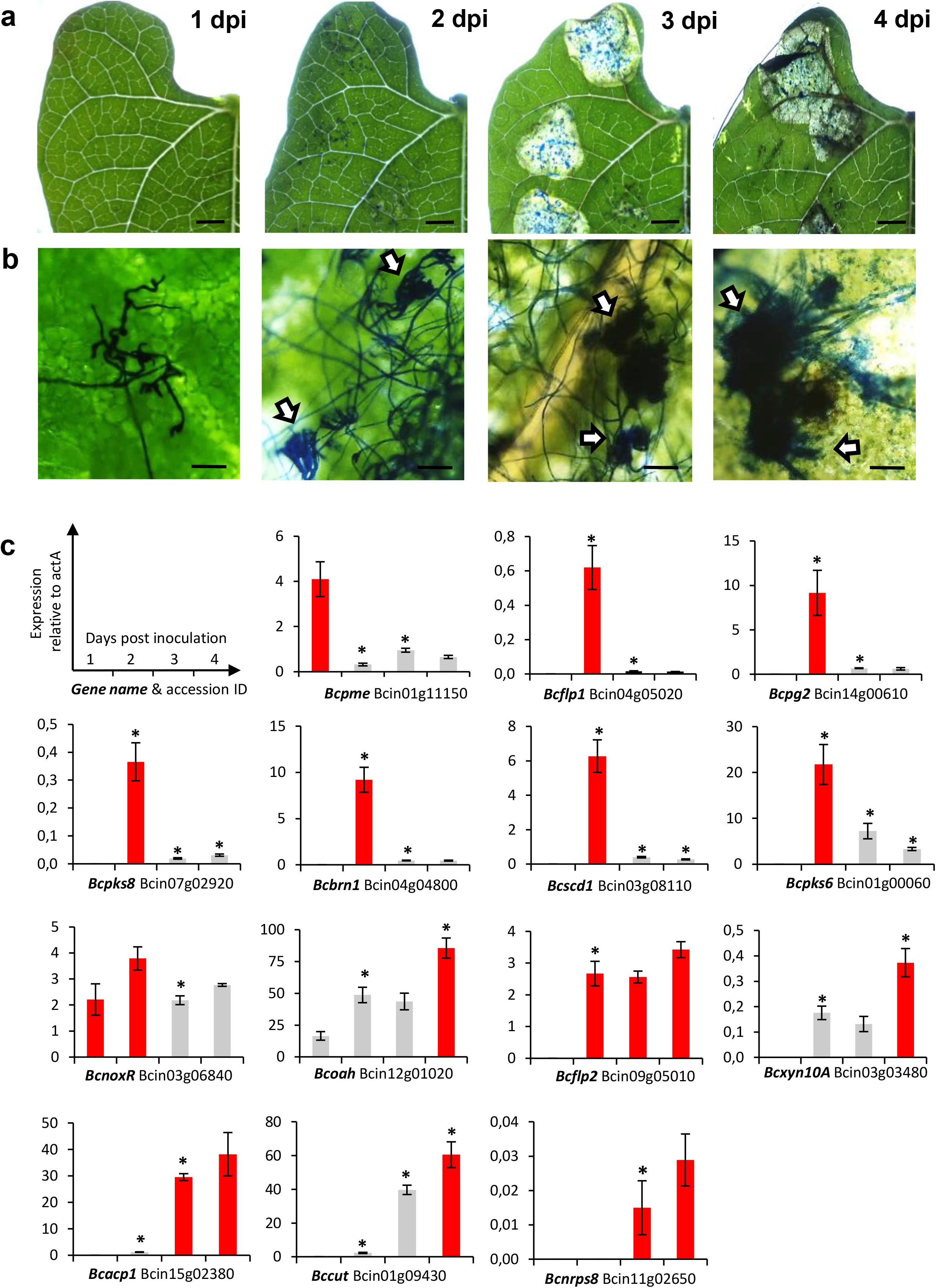
*In planta* RT-qPCR validation of microarray up-regulated genes. (a) and (b) Infection cushions development on leaves infected by *B. cinerea*. Primary bean leaves were inoculated with conidia and fungal development (1-4 days post inoculation (dpi)) was monitored using a Stereomicroscope (Zeiss) and cotton blue to stain the fungal cells at the plant surface. IC are pointed by white arrows. Bars represent 5 mm (a) or 50 μm (b). (c) Expression of selected genes during infection. Infected bean leaves were collected to prepare RNAs and RT-qPCR was used to measure the expression of 14 genes up-regulated in IC produced *in vitro* conditions (microarray data) and whose predicted function relates to virulence. The red bar indicates the peak of expression for each gene (when 2 or 3 bars are in red for the same gene, statistics cannot distinguish them). Gene expression levels were calculated following the 2^−ΔΔ*C*T^ method using constitutively expressed actin gene *BcactA* (Bcin16g02020) as a reference. The use of two other housekeeping genes, elongation factor *Bcef1α* (Bcin09g05760) and pyruvate dehydrogenase *Bcpda1* (Bcin07g01890) showed similar results (data not shown). Standard errors of three independent biological replicates are displayed and asterisks indicate a significant difference in gene expression compared with the previous time point (Student’s *t* test, p-value <0.05).

### Two fasciclin genes up-regulated in IC are virulence factors in *B. cinerea*

Two genes encoding fasciclin-like proteins (*Bcflp1*; Bcin04g05020 and *Bcflp2*; Bcin09g05010) were selected for mutagenesis. Fasciclins in fungi could play a role in development or in autophagy (Seifert, 2018) and their role in the virulence of the rice pathogen *M. oryzae* has been shown (Liu *et al.*, 2009). The *Bcflp1* and *Bcflp2* genes were both up-regulated in IC and this differential expression was confirmed by RT-qPCR *in vitro* and *in planta* (Table S2; Fig. 5). Besides, BcFlp1 was up-accumulated in the secretome of IC (Table S3). By using a gene replacement strategy, two single mutant strains (*ΔBcflp1* and *ΔBcflp2*) and the double mutant strain (*ΔBcflp1::ΔBcflp2*) were constructed. These strains were genetically purified and their genotype were verified by PCR and Southern blotting (Fig. S5). The formation of IC could be observed in all mutants (data not shown) and this result indicates that the two fasciclin-like proteins are not required for the differentiation of IC in *B. cinerea.* Deletion of *Bcflp1* did not alter the hyphal growth *in vitro* (Fig. 6a) but it significantly delayed the colonization phase on bean leaves (Fig. 6b-c). Deletion of *Bcflp2* caused a 20% growth reduction on minimal medium *in vitro* (Fig. 6a). It also caused a reduction and then an arrest of bean leaves colonization at 7 dpi (data not shown) while a dark ring could be observed at the periphery of the infected zone (Fig. 6b-c). At last, the phenotype of the *ΔBcflp1::ΔBcflp2* mutant was similar to that of the *ΔBcflp2* mutant (Fig. 6).

**Figure 6:**
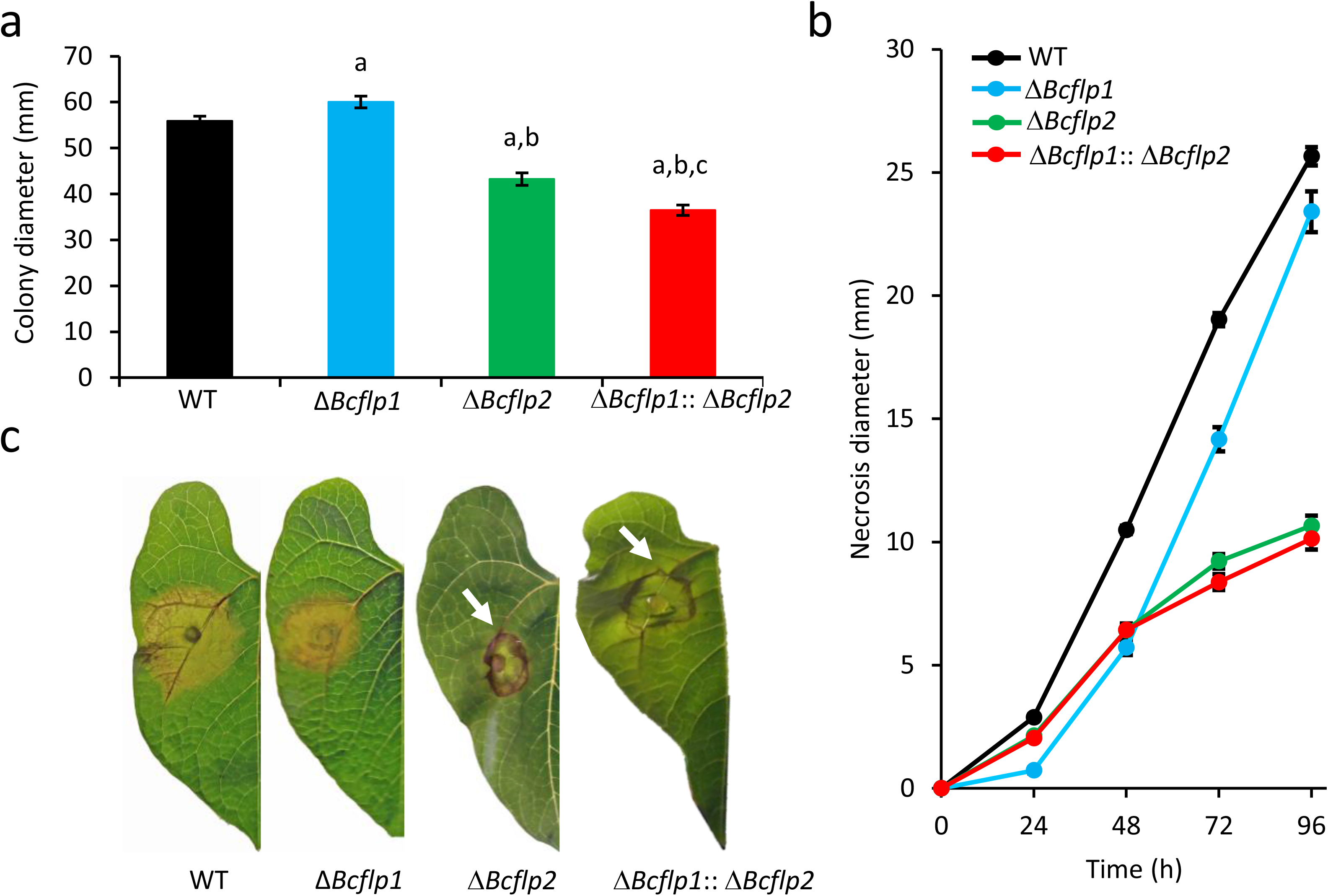
Characterization of fasciclin-like deletion mutants in *Botrytis cinerea*. (a) Growth *in vitro*: The WT strain, the single mutant strains (Δ*Bcflp1* and Δ*Bcflp2*) and the double mutant strain (Δ*Bcflp1*::Δ*Bcflp2*) were grown on minimal medium for 72h. The diameters of the colonies were measured on 4 independent experiments conducted with 5 plates for each strain. Bars indicate standard errors. Letters indicate significant differences (Student *t*-test, p-value <0.05) between: (a) WT and mutants, (b) Δ*Bcflp1* and Δ*Bcflp2* or Δ*Bcflp1*::Δ*Bcflp2*, (c) Δ*Bcflp2* and Δ*Bcflp1*::Δ*Bcflp2*. (b) Infection kinetics - French bean leaves were inoculated with mycelial plugs of the WT and deletion strains. The necrosis of the plant tissues was measured every 24 h (at least eight plants and 32 infection points) over 4 days in 3 independent experiments. Bars indicate standard errors. (c) Visualization of the necrotic lesions. At 72 h post inoculation, typical images of bean leaves infected by the WT and mutant strains were recorded. White arrows indicate dark rings at the edge of necrotic zones. Several clones were obtained for each deletion strain and tested for virulence. As they behaved similarly, results are shown for only one clone.

## Discussion

In plant pathogenic fungi, IC have been proposed to play an important role in host penetration. In this study, four cutinase-encoding genes and 48 predicted PCWDE-encoding genes were revealed as up-regulated in *B. cinerea* IC. These genes include the endopolygalacturonase *BcPg2* (Bcin14g00610), the endoxylanase *BcXyn11A* (Bcin03g00480), and the endoarabinanase *BcAra1* (Bcin02g07700) required for full virulence of *B. cinerea* (Kars *et al*., 2005; Brito *et al*., 2006; Nafisi *et al*., 2014). Besides, aspartyl proteases, sedolisins and metallopeptidases-encoding genes were also significantly up-regulated. A proteomic analysis of IC secretome confirmed the up-accumulation of CAZymes and proteases when compared to vegetative mycelium, and altogether these results support a role for IC in host penetration *via* enzymatic degradation of plant cell walls.

Interestingly, genes coding for membrane transporters and catabolic pathways were up-regulated in IC. The up-regulation of sugar-transporters suggests an increase in sugar uptake inside IC while the up-regulation of metabolic pathways involved in glucose metabolism suggests a nutritional role for IC. In connection with the up-regulation of PCWDE-encoding genes, IC could be considered as fungal structures equipped with the necessary tools to recycle the plant cell wall components and/or to feed upon the products of plant polysaccharides degradation.

The transcriptomic data highlighted IC as structures also dedicated to the establishment of the necrotrophic lifestyle of *B. cinerea* through the secretion of phytotoxins, ROS, and necrosis inducers. Indeed, two known SM gene clusters are up-regulated in IC. These clusters are responsible for the production of botcinic acid and botrydial, two phytotoxins playing a role in plant infection by *B. cinerea* (Cutler *et al.*, 1993 & 1996; Deighton *et al.*, 2001; Dalmais *et al.*, 2011; Massaroli *et al*., 2013; Collado & Viaud, 2016). Botrydial sesquiterpene and its derivatives botryanes are proposed as fungal effectors manipulating plant host defenses, promoting cell death and thus enabling *B. cinerea* to feed on necrotic tissues (Rossi *et al*., 2011). Botcinic acid polyketide and its structurally related botcinins and botrylactones cause plant chlorosis and necrosis (Cutler *et al*., 1993). Double inactivation of these SM clusters led to a defect of pathogenesis and demonstrated the concerted action of the two toxins (Dalmais *et al*., 2011). Interestingly, the co-expression of these clusters in IC supports this concerted action. In addition, four putative SM clusters were also up-regulated in IC. These clusters might play a role in virulence, but this and the putative metabolites produced as a result of their activation await characterization. It is noteworthy that several orphan SM have been reported in *B. cinerea* while their biosynthesis genes remain unknown (Collado and Viaud, 2016). Additionally, it was proposed that secretion of new sesquiterpenoid metabolites called eremophilenols could promote and regulate the production of *B. cinerea* IC, perhaps acting as an endogenous signal (Pinedo *et al*., 2016).

During the early phase of plant-fungal interaction, ROS are produced by the plants, as part of their defense mechanisms, but they are also produced by the pathogen as cell death inducers. In order to produce and to cope with ROS, *B. cinerea* is equipped with multiple oxidoreductases. Chemical compounds or mutations that target some of these enzymes, or their encoding genes, impair virulence (Rolke *et al.*, 2004; Segmüller *et al.*, 2008; An *et al.*, 2016; Siegmund & Viefhues, 2016). The analysis of the up-regulated genes in IC highlighted 136 genes related to the oxidation-reduction process, among which 15 genes coding for putative ROS producing enzymes. In parallel, relatively higher levels of H2O2 were also detected in the IC culture medium compared to control vegetative mycelium. This suggests that IC secretes high amounts of ROS, which would be consistent with the production of ROS observed in IC during penetration events of onion epidermis (Choquer *et al*., 2007; Marschall & Tudzynski, 2016b). Several genes have already been identified in recent years whose mutation has a negative impact on IC formation in *B. cinerea*. Interestingly these genes are often associated to the oxidation-reduction process and play a role in the development and virulence of *B. cinerea*. These genes encode the regulator of oxidative stress response BcSkn7 (Viefhues *et al*., 2015), the scaffold protein BcIqg1 involved in resistance against oxidative stress (Marschall and Tudzynski, 2016a), the aquaporin BcAqp8 involved in ROS production (An *et al*., 2016), the ER protein BcPdi1 involved in redox homeostasis (Marschall and Tudzynski, 2017), the septin BcSep4 whose proper assembly is regulated by ROS (Feng *et al*., 2017), and the H3K4 demethylase BcJar1 orchestrating ROS production (Hou *et al*., 2020). It has been proposed that ROS are required for the formation and functioning of UA (Ryder and Talbot, 2015), and several recent publications as well as this study seem to corroborate the essential role of ROS in the development of IC.

*B. cinerea* secretes necrosis-inducing factors during the early phase of plant infection (Shlezinger *et al.*, 2011; González *et al.*, 2016). This would trigger a hypersensitive response (HR) from which the fungus would benefit to achieve full virulence (Govrin and Levine, 2000; Govrin *et al.*, 2006). Such factors are exemplified by BcNEPs (Qutob *et al.*, 2007; Arenas *et al.*, 2010) and both the transcriptomic and proteomic data of IC revealed the *Bcnep2* gene and the Bcnep2 protein as up-regulated and up-accumulated, respectively. Similar results were obtained for the xyloglucanase BcXyg1 and the xylanase Bcxyn11A, two additional necrosis inducers (Noda *et al*., 2010; Zhu *et al*., 2017). Together with the up-regulation of genes involved in the production of phytotoxins and ROS, this argues for IC playing a role in the killing of plant cells.

In *B. cinerea*, IC cell walls are darkened by DHN-melanins. These pigments are secondary metabolites produced via a bipartite pathway in *B. cinerea*, so far identified as specific to conidia or sclerotia (Schumacher, 2016). The gene expression data reported in this study indicate that the conidial metabolic pathway is activated in IC while the sclerotial pathway is not. This result suggested that IC produce more melanin than vegetative hyphae. Microscopic observation of thick dark cell walls in IC that are sensitive to tricyclazole inhibitor of DHN-melanins supported this hypothesis. Melanins biosynthesis is dispensable for virulence in *B. cinerea* (Schumacher, 2016) and the reason of their increased production in IC remains to be clarified. It could play a role in the fungus survival during the oxidative burst of host plants (Govrin and Levine, 2000). In other fungi, melanins were proposed to play a role in the crosslinking of cell wall compounds (Franzen *et al.*, 2006), the binding of proteins (Mani *et al.*, 2001) or that of metals (Fogarty and Tobin, 1996).

Three genes coding for chitin deacetylases (CDA) were also up-regulated in IC when compared to vegetative hyphae. This suggests that chitin is deacetylated into chitosan in the cell wall of IC and differential staining of these two compounds allowed microscopic observations that brought experimental support to this hypothesis. Since CDA can play a role in polar growth of *Aspergillus fumigatus* (Xie *et al.*, 2020) or in appressorium differentiation of *Magnaporthe oryzae* (Kuroki *et al.*, 2017), the transformation of chitin into chitosan in *B. cinerea* could be involved in the change of hyphal growth that leads to IC formation. The production of chitosan by CDA could also play a role in cell wall anchoring of melanins, cell wall integrity, osmotic stability, modulation of extra-cellular polysaccharide production and/or adhesion to surfaces (Baker *et al.*, 2007; Geoghegan and Gurr, 2016; Perez-Dulzaides *et al.*, 2018). Indeed, the role of chitosan as a critical component of the fungal cell-wall scaffold governing DOPA-melanin deposition was verified in Cryptococcal species (Chrissian *et al*., 2020). At last, deacetylation of chitin in plant endophytic fungi can prevent detection of chitin oligomers by the host immune system (Cord-Landwehr *et al.*, 2016) and whether a similar process of camouflage would exist in IC of *B. cinerea* could also be considered. Indeed, chitosan biosynthesis was found to be a critical factor for the virulence of Cryptococcal species (Upadhya *et al*., 2018; Lam *et al*., 2019). Among the other up-regulated fungal cell wall enzymes, we can cite the beta-glucosidase BcSun1 (Bcin06g06040; Pérez-Hernández *et al*., 2017) that is required for the full production of IC and the full virulence of *B. cinerea*.

A two-member family of fascicline-like proteins was selected for mutagenesis based on the up-regulation of their encoding genes and up-accumulation of one of them in the secretome of IC. The delayed or impaired leaves colonization that was observed in the mutant strains suggests a role for these proteins at a later stage of the infectious process. As fasciclins could be involved in autophagy in *M. oryzae* and *Schizosaccharomyces pombe* (Liu *et al*., 2009; Sun *et al*., 2013), one may hypothesize a similar function for BcFlp1 and/or BcFlp2, but this needs to be explored.

In a recent study, we have studied mutants of *B. cinerea* that do not produce IC and have collected transcriptomic and proteomic data for four of them (De Vallée *et al.*, 2019). Comparison of the microarray data presented in this study to those collected for these mutants showed that 40 of the first 100 up-regulated genes in IC are down-regulated in all 4 mutants (Fig. S6). This inverted correlation was strengthened by the comparison of the secretomes since 47% of the proteins up-accumulated in the culture medium of IC were down-accumulated in the 4 IC-deficient mutants (Table S3). These genes represent a list of putative targets for mutagenesis and functional studies that could reveal their importance in the pathogenicity of *B. cinerea*.

In conclusion, the transcriptomic study of mature IC of *B. cinerea* provided information that support an involvement of these structures in intoxication of plant cells, plant material degradation and possibly fungal nutrition. These responses were also observed in the transcriptomes of *B. cinerea* infecting plant hosts such as lettuce, tomato, grapevine, cucumber or *Arabidopsis* (Blanco-Ulate *et al*., 2014; Kelloniemi *et al*., 2015; Kong *et al*., 2015; Zhang *et al*., 2019). Although IC were produced *in vitro*, this study revealed that even in the absence of contact with plants, PCWDE and cutinase encoding genes were still upregulated by the fungus. This suggests that IC formation and activation of genes involved in downstream events could be triggered by the perception of a single signal, possibly that of a hard surface. Could IC be a “weapon” allowing the fungus to trigger plant disease? The recent evidence that several apathogenic mutants are no longer able to produce normal IC would support this hypothesis.

## Acknowledgements

We thank Adeline Simon, Lucile Albinet-Mauprivez and Luna Nadjare for providing tools and help on bioinformatic analyses. We thank Gwenlyn Fleury for her implication in the study of fasclicin mutants, and Cindy Dieryckx and Vincent Girard for their advices on secretome analysis.

## Author contributions (by alphabetical order)

Original concept (CN, JAR, MC, NP); Writing the manuscript (CB, CN, CRa, MC, NP); Editing the manuscript (CB, CN, CRa, IRG, MC, NP); Microarrays experiment (CRa); Transcriptome bioinformatics (CB, CRa, EL, IRG, MC); Fasciclin mutants construction and phenotyping (CRa, JF, MC, PS); Proteomic experiment and analysis(CRi, JWD); Microscopy (AdV, CB, CRa, ES, JAR, NP, RM); qPCR (CRa, MJG; MS).

## Supporting Information

**Figure S1:** *In vitro* production of infection cushions of *Botrytis cinerea*.

**Figure S2:** Gene ontology (GO) term enrichment analysis of genes differentially expressed in the infection cushion of *Botrytis cinerea*.

**Figure S3:** Regulation of secondary metabolism biosynthesis key genes in the infection cushion of *Botrytis cinerea.*

**Figure S4:** Putative secondary metabolism gene clusters up-regulated in the infection cushion of *Botrytis cinerea.*

**Figure S5:** Mutagenesis of genes encoding fasciclin-like proteins in *Botrytis cinerea*.

**Figure S6:** Comparison of the *Botrytis cinerea* infection cushion (IC) transcriptome and the transcriptomes of four IC-deficient mutants.

**Table S1:** Results of the microarray analysis of the infection cushion of *Botrytis cinerea*.

**Table S2:** RT-qPCR validation of microarray expression profiles from the infection cushion of *Botrytis cinerea* produced *in vitro*.

**Table S3:** Up-accumulated proteins in the secretome of the infection cushion of *Botrytis cinerea.*

**Table S4:** Constructs and primers used in this study.

## References

Akutsu K, Kobayashi Y, Matsuzawa Y, Watanabe T, Ko K, Misato T. 1981. Morphological studies on infection process of cucumber leaves by conidia of *Botrytis cinerea* stimulated with various purine-related compounds. Annals of the Phytopathological Society of Japan 47: 234–243.

Almagro Armenteros JJ, Tsirigos KD, Sønderby CK, Petersen TN, Winther O, Brunak S, von Heijne G, Nielsen H. 2019. SignalP 5.0 improves signal peptide predictions using deep neural networks. Nature Biotechnology 37: 420–423.

Amselem J, Cuomo CA, van Kan JA, Viaud M, Benito EP, Couloux A, Coutinho PM, deVries RP, Dyer PS, Fillinger S et al. 2011. Genomic analysis of the necrotrophic fungal pathogens *Sclerotinia sclerotiorum* and *Botrytis cinerea*. PLoS Genetics 7: e1002230.

An B, Li B, Li H, Zhang Z, Qin G, Tian S. 2016. Aquaporin8 regulates cellular development and reactive oxygen species production, a critical component of virulence in *Botrytis cinerea*. New Phytologist 209: 1668–1680.

Arenas YC, Kalkman E, Schouten A, Dieho M, Vredenbregt P, Uwumukiza B, Ruiz MO, van Kan JA. 2010. Functional analysis and mode of action of phytotoxic Nep1-like proteins of *Botrytis cinerea*. Physiological and Molecular Plant Pathology 74: 376–386.

Backhouse D, Willets HJ. 1987. Development and structure of infection cushions of *Botrytis cinerea*. Transactions of the British Mycological Society 89: 89–95.

Baker LG, Specht CA, Donlin MJ, Lodge JK. 2007. Chitosan, the deacetylated form of chitin, is necessary for cell wall integrity in *Cryptococcus neoformans*. Eukaryotic Cell 6: 855–867.

Blanco-Ulate B, Morales-Cruz A, Amrine KC, Labavitch JM, Powell AL, Cantu D. 2014. Genome-wide transcriptional profiling of *Botrytis cinerea* genes targeting plant cell walls during infections of different hosts. Frontiers in Plant Science 5: 435.

Boenisch MJ, Schäfer W. 2011. *Fusarium graminearum* forms mycotoxin producing infection structures on wheat BMC Plant Biology 11:110.

Brito N, Espino JJ, González C. 2006. The endo-β-1, 4-xylanase Xyn11A is required for virulence in *Botrytis cinerea*. Molecular Plant-Microbe Interactions 19: 25–32.

Catlett NL, Lee B-N, Yoder OC, Turgeon BG. 2003. Split-Marker Recombination for Efficient Targeted Deletion of Fungal Genes. Fungal Genetics Reports 50: 9–11.

Choquer M, Fournier E, Kunz C, Levis C, Pradier JM, Simon A, Viaud M. 2007. *Botrytis cinerea* virulence factors: new insights into a necrotrophic and polyphageous pathogen. FEMS Microbiology Letters 277:1–10.

Chrissian C, Camacho E, Shun Fu M, Prados-Rosales R, Chatterjee S, Cordero RJB, Lodge JK, Casadevall A, Stark RE. 2020. Melanin deposition in two *Cryptococcus* species depends on cell-wall composition and flexibility. Journal of Biological Chemistry 295: 1815–1828.

Cohrs KC, Simon A, Viaud M, Schumacher J. 2016. Light governs asexual differentiation in the grey mould fungus *Botrytis cinerea* via the putative transcription factor BcLTF2. Environmental Microbiology 18: 4068–4086.

Collado IG, Viaud M. 2016. Secondary metabolism in *Botrytis cinerea*: combining genomic and metabolomic approaches. In: Fillinger S, Elad Y, eds. Botrytis-the fungus, the pathogen and its management in agricultural systems. Cham, Switzerland: Springer: 291–313.

Cord-Landwehr S, Melcher RLJ, Kolkenbrock S, Moerschbacher BM. 2016. A chitin deacetylase from the endophytic fungus *Pestalotiopsis* sp efficiently inactivates the elicitor activity of chitin oligomers in rice cells. Scientific Reports 6: 38018.

Cutler HG, Jacyno JM, Harwood JS, Dulik D, Goodrich PD, Roberts RG. 1993. Botcinolide: A Biologically Active Natural Product from *Botrytis cinerea*. Bioscience, Biotechnology, and Biochemistry 57:1980–1982.

Cutler HG, Parker SR, Ross SA, Crumley FG, Schreiner PR. 1996. Homobotcinolide: A biologically active natural homolog of botcinolide from *Botrytis cinerea*. Bioscience, Biotechnology, and Biochemistry 60: 656–658.

Dalmais B, Schumacher J, Moraga J, Le Pecheur P, Tudzynski B, Collado IG, Viaud M. 2011. The *Botrytis cinerea* phytotoxin botcinic acid requires two polyketide synthases for production and has a redundant role in virulence with botrydial. Molecular Plant Pathology 12: 564–579.

Daniels A, Lucas JA, Peberdy JF. 1991. Morphology and ultrastructure of W and R pathotypes of *Pseudocercosporella herpotrichoides* on wheat seedlings. Mycological Research 95: 385–397.

De Vallée A, Bally P, Bruel C, Chandat L, Choquer M, Dieryckx C, Dupuy JW, Kaiser S, Latorse MP, Loisel E et al. 2019. A similar secretome disturbance as a hallmark of non-pathogenic *Botrytis cinerea* ATMT-mutants? Frontiers in Microbiology 10: 2829.

Dean R, van Kan JA, Pretorius ZA, Hammond-Kosack KE, Di Pietro A, Spanu PD, Rudd JJ, Dickman M, Kahmann R, Ellis J et al. 2012. The top 10 fungal pathogens in molecular plant pathology. Molecular Plant Pathology 13: 414–430.

Deighton N, Muckenschnabel I, Colmenares AJ, Collado IG, Williamson, B. 2001. Botrydial is produced in plant tissues infected by *Botrytis cinerea*. Phytochemistry 57: 689–692.

Deising HB, Werner S, Wernitz M. 2000. The role of fungal appressoria in plant infection. Microbes and Infection 2: 1631–41.

Demirci E, Döken MT. 1998. Host Penetration and Infection by the anastomosis groups of *Rhizoctonia solani* Kühn isolated from potatoes. Turkish Journal of Agriculture and Forestry 22: 609–613.

Demoor A, Silar P, Brun S. 2019. Appressorium: The Breakthrough in *Dikarya*. Journal of Fungi 5: 72.

Dieryckx C, Gaudin V, Dupuy JW, Bonneu M, Girard V, Job D. 2015. Beyond plant defense: insights on the potential of salicylic and methylsalicylic acid to contain growth of the phytopathogen *Botrytis cinerea*. Frontiers in Plant Science 6: 859.

Dinh SQ, Joyce DC, Irving DE, Wearing AH. 2011. Histology of waxflower (*Chamelaucium spp*.) flower infection by *Botrytis cinerea*. Plant Pathology 60: 278–287.

Elad Y, Pertot I, Marina A, Prado AM, Stewart A. 2016. Plant hosts of Botrytis spp. In: Fillinger S, Elad Y, eds. Botrytis-the fungus, the pathogen and its management in agricultural systems. Cham, Switzerland: Springer: 413–486.

Emmett RW, Parbery DG 1975. Appressoria. Annual Review of Phytopathology 13: 147–165.

Feng HQ, Li GH, Du SW, Yang S, Li XQ, de Figueiredo P, Qin QM. 2017. The septin protein Sep4 facilitates host infection by plant fungal pathogens via mediating initiation of infection structure formation. Environmental Microbiology 19: 1730–1749.

Fogarty RV, Tobin JM. 1996. Fungal melanins and their interactions with metals. Enzyme and Microbial Technology 19: 311–317.

Fourie JF, Holz G. 1994. Infection of plum and nectarine flowers by *Botrytis cinerea*. Plant Pathology 43: 309–315.

Franzen AJ, Cunha MM, Batista EJ, Seabra SH, De Souza W, Rozental S. 2006. Effects of tricyclazole 5-methyl-1,2,4-triazol[3,4] benzothiazole, a specific DHN-melanin inhibitor, on the morphology of *Fonsecaea pedrosoi* conidia and sclerotic cells. Microscopy Research and Technique 69: 729–737.

Frias M, Brito N, González C. 2013. The *Botrytis cinerea* cerato‐platanin BcSpl1 is a potent inducer of systemic acquired resistance (SAR) in tobacco and generates a wave of salicylic acid expanding from the site of application. Molecular Plant Pathology 14: 191–196.

Frias M, Gonzalez M, Gonzalez C, Brito N. 2016. BcIEB1, a *Botrytis cinerea* secreted protein, elicits a defense response in plants. Plant Science 250: 115–124.

Fullerton RA, Harris FM, Hallett IC. 1999. Rind distortion of lemon caused by *Botrytis cinerea* Pers. New Zealand Journal of Crop and Horticultural Science 27: 205–214.

Garcia-Arenal F, Sagasta EM. 1980. Scanning electron microscopy *of Botrytis cinerea* penetration of bean *Phaseolus vulgaris* hypocotyls. Journal of Phytopathology 99: 37–42.

Geoghegan IA, Gurr SJ. 2016. Chitosan mediates germling adhesion in *Magnaporthe oryzae* and is required for surface sensing and germling morphogenesis. PLoS Pathogens 126: e1005703.

Gladders P, Coley-Smith JR. 1977. Infection cushion formation in Rhizoctonia tuliparum, Transactions of the British Mycological Society 68: 115–118.

Glass NL, Schmoll M, Cate JH, Coradetti S. 2013. Plant cell wall deconstruction by ascomycete fungi. Annual Review of Microbiology 67: 477–498.

González C, Brito N, Sharon A. 2016. Infection Process and Fungal Virulence Factors. In: Fillinger S, Elad Y, eds. Botrytis-the fungus, the pathogen and its management in agricultural systems. Cham, Switzerland: Springer: 229–246.

Govrin EM, Levine A. 2000. The hypersensitive response facilitates plant infection by the necrotrophic pathogen *Botrytis cinerea*. Current Biology 10: 751–757.

Govrin EM, Rachmilevitch S, Tiwari BS, Solomon M, Levine A. 2006. An elicitor from *Botrytis cinerea* induces the hypersensitive response in *Arabidopsis thaliana* and other plants and promotes the gray mold disease. Phytopathology 96: 299–307.

Hao FM, Ding T, Wu MD, Zhang J, Yang L, Chen WD, Li GQ. 2018. Two novel hypovirulence-associated mycoviruses in the phytopathogenic fungus *Botrytis cinerea*: molecular characterization and suppression of infection cushion formation. Viruses 10: 254.

Hou J, Feng H‐Q, Chang H‐W, Liu Y, Li G‐H, Yang S, Sun C‐H, Zhang M‐Z, Yuan Y, Sun J, et al. 2020. The H3K4 demethylase Jar1 orchestrates ROS production and expression of pathogenesis‐related genes to facilitate *Botrytis cinerea* virulence. New Phytologist 225: 930–947.

Käll L, Canterbury J, Weston J, Noble WS, MacCoss MJ. 2007. Semi-supervised learning for peptide identification from shotgun proteomics datasets, Nature Methods 4: 923–925.

Kars I, Krooshof GH, Wagemakers L, Joosten R, Benen JA, van Kan JA. 2005. Necrotizing activity of five *Botrytis cinerea* endopolygalacturonases produced in *Pichia pastoris*. Plant Journal 43: 213–25.

Kelloniemi J, Trouvelot S, Heloir MC, Simon A, Dalmais B, Frettinger P, Cimerman A, Fermaud M, Roudet J, Baulande S et al. 2015. Analysis of the Molecular Dialogue Between Gray Mold (*Botrytis cinerea*) and Grapevine (*Vitis vinifera*) Reveals a Clear Shift in Defense Mechanisms During Berry Ripening. Molecular Plant-Microbe Interactions 28: 1167–1180.

Kong WW, Chen N, Liu TT, Zhu J, Wang JQ, He XQ, Jin Y. 2015. Large-Scale Transcriptome Analysis of Cucumber and *Botrytis cinerea* during Infection. PLoS One 10: e0142221.

Kuroki M, Okauchi K, Yoshida S, Ohno Y, Murata S, Nakajima Y, Nozaka A, Tanaka N, Nakajima M, Taguchi H et al. 2017. Chitin-deacetylase activity induces appressorium differentiation in the rice blast fungus *Magnaporthe oryzae*. Scientific Reports 7: 9697.

Lalève A, Gamet S, Walker A‐S, Debieu D, Toquin V, Fillinger S. 2014. Roles of key SdhB residues in SDH activity and inhibition. Environmental Microbiology 16: 2253–2266.

Lam WC, Upadhya R, Specht CA, Ragsdale AE, Hole CR, Levitz SM, Lodge JK. 2019. Chitosan biosynthesis and virulence in the human fungal pathogen *Cryptococcus gattii*. mSphere 4: e00644–19.

Liu TB, Chen GQ, Min H, Lin FC. 2009. MoFlp1, encoding a novel fungal fasciclin-like protein, is involved in conidiation and pathogenicity in *Magnaporthe oryzae*. Journal of Zhejiang University Science B 10: 434–444.

Liu Z, Gay LM, Tuveng TR, Agger JW, Westereng B, Mathiesen G, Horn SJ, Vaaje-Kolstad G, van Aalten DM, Eijsink VG. 2017. Structure and function of a broad-specificity chitin deacetylase from *Aspergillus nidulans* FGSC A4. Scientific Reports 7: 1746.

Liu JK, Chang HW, Liu Y, Qin YH, Ding YH, Wang L, Zhao Y, Zhang MZ, Cao SN, Li LT et al. 2018. The key gluconeogenic gene PCK1 is crucial for virulence of *Botrytis cinerea* via initiating its conidial germination and host penetration. Environmental Microbiology 20: 1794–1814.

Livak KJ, Schmittgen TD. 2001. Analysis of relative gene expression data using real-time quantitative PCR and the 2(-ΔΔC(T)) method. Methods 25: 402–408.

Lombard V, Golaconda Ramulu H, Drula E, Coutinho PM, Henrissat B. 2013. The carbohydrate-active enzymes database (CAZy) in 2013. Nucleic Acids Research 42: D490–D495.

Lumsden RD, Wergin WP. 1980. Scanning-Electron Microscopy of Infection of Bean by Species of *Sclerotinia*. Mycologia 72: 1200–1209.

Mani I, Sharma V, Tamboli I, Raman G. 2001. Interaction of melanin with proteins - the importance of an acidic intramelanosomal pH. Pigment Cell Research 14: 170–179.

Marschall R, Tudzynski P. 2016a. BcIqg1, a fungal IQGAP homolog, interacts with NADPH oxidase, MAP kinase and calcium signaling proteins and regulates virulence and development in *Botrytis cinerea*. Molecular Microbiology 101: 281–298.

Marschall R, Tudzynski P. 2016b. Reactive oxygen species in development and infection processes. Seminars in Cell & Developmental Biology 57: 138–146.

Marschall R, Tudzynski P. 2017. The Protein disulfide isomerase of *Botrytis cinerea*: An ER protein involved in protein folding and redox homeostasis influences NADPH oxidase signaling processes. Frontiers in Microbiology 8: 960.

Massaroli M, Moraga J, Borges KB, Ramirez-Fernandez J, Viaud M, Collado IG, Duran-Patron R, Hernandez-Galan R. 2013. A Shared Biosynthetic Pathway for Botcinins and Botrylactones Revealed through Gene Deletions. ChemBioChem 14: 132–136.

Michielse CB, Becker M, Heller J, Moraga J, Collado G, Tudzynski P. 2011. The *Botrytis cinerea* Reg1 protein, a putative transcriptional regulator, is required for pathogenicity, co-nidiogenesis, and the production of secondary metabolites. Molecular Plant-Microbe Interactions 24: 1074–1085.

Mitchell AL, Attwood TK, Babbitt PC, Blum M, Bork P, Bridge A, Brown SD, Chang HY, El-Gebali S, Fraser MI et al. 2019. InterPro in 2019: improving coverage, classification and access to protein sequence annotations. Nucleic Acids Research 47: D351–D360.

Nafisi M, Stranne M, Zhang L, van Kan JA, Sakuragi Y. 2014. The Endo-Arabinanase BcAra1 Is a Novel Host-Specific Virulence Factor of the Necrotic Fungal Phytopathogen *Botrytis cinerea*. Molecular Plant Microbe Interactions 27: 781–92

Noda J, Brito N, González C. 2010. The *Botrytis cinerea* xylanase Xyn11A contributes to virulence with its necrotizing activity, not with its catalytic activity. BMC Plant Biology 10: 38.

Pepin R. 1980. Le comportement parasitaire de *Sclerotinia tuberosa* (hedw.) Fuckel sur *Anemone nemorosa* L. Etude en microscopie photonique et electronique à balayage. Mycopathologia 72: 89.

Perez-Dulzaides R, Camacho E, Cordero RJB, Casadevall A. 2018. Cell-wall dyes interfere with *Cryptococcus neoformans* melanin deposition. Microbiology 164: 1012–1022.

Perez-Hernandez A, Gonzalez M, Gonzales C, van Kan JA, Brito N. 2017. BcSUN1, a *B.cinerea* SUN-family protein is involved in Virulence. Frontiers in Microbiology 8: 35.

Perez-Riverol Y, Csordas A, Bai J, Bernal-Llinares M, Hewapathirana S, Kundu DJ, Inuganti A, Griss J, Mayer G, Eisenacher M et al. 2019. The PRIDE database and related tools and resources in 2019: improving support for quantification data. Nucleic Acids Research 47: D442–D450.

Pineda E, Thonnus M, Mazet M, Mourier A, Cahoreau E, Kulyk H, Dupuy JW, Biran M, Masante C, Allmann S et al. 2018. Glycerol supports growth of the *Trypanosoma brucei* bloodstream forms in the absence of glucose: Analysis of metabolic adaptations on glycerol-rich conditions. PLoS Pathogens 14: e1007412.

Pinedo C, Moraga J, Barua J, González-Rodríguez VE, Aleu J, Durán-Patrón R, Mací-as-Sánchez AJ, Hanson JR, Viaud M, Hernández-Galán R. 2016. Chemically induced cryptic sesquiterpenoids and expression of sesquiterpene cyclases in *Botrytis cinerea* revealed new sporogenic (+)-4-Epi eremophil-9-en-11-ols. ACS Chemical Biology 11: 1391–1400.

Porquier A, Moraga J, Morgant G, Dalmais B, Simon A, Sghyer H, Collado IG,Viaud M. 2019. Botcinic acid biosynthesis in *Botrytis cinerea* relies on a subtelomeric gene cluster surrounded by relics of transposons and is regulated by the Zn2-Cys6 transcription factor BcBoa13. Current Genetics 65: 965–980.

Priebe S, Kreisel C, Horn F, Guthke R, Linde J. 2015. FungiFun2: a comprehensive online resource for systematic analysis of gene lists from fungal species. Bioinformatics 31: 445–446.

Prior GD, Owen JH. 1964. Pathological anatomy of *Sclerotinia trifoliorum* on clover and alfalfa. Phytopathology 54:784–787.

Qutob D, Kemmerling B, Brunner F, Küfner I, Engelhardt S, Gust A, Luberacki B, Seitz H, Stahl D, Rauhut, T. 2007. Phytotoxicity and innate immune responses induced by Nep1-like proteins. Plant Cell 18: 3721–3744.

Rascle C, Dieryckx C, Dupuy JW, Muszkieta L, Souibgui E, Droux M, Bruel C, Girard V, Poussereau N. 2018. The pH regulator PacC: a host-dependent virulence factor in *Botrytis cinerea*. Environmental Microbiology Reports 10: 555–568.

Rheinländer PA, Sutherland PW, Fullerton RA. 2013. Fruit infection and disease cycle of *Botrytis cinerea* causing cosmetic scarring in persimmon fruit (*Diospyros kaki Linn*.). Australasian Plant Pathology 42: 551–560.

R Core Team. 2013. R: A language and environment for statistical computing. Vienna, Austria: R Foundation for Statistical Computing. [WWWdocument] URL http://www.R-project.org/

Rolke Y, Liu SJ, Quidde T, Williamson B, Schouten A, Weltring KM, Siewers V, Tenberge KB, Tudzynski B, Tudzynski P. 2004. Functional analysis of H_2_O_2_-generating systems in *Botrytis cinerea*: the major Cu-Zn-superoxide dismutase (BCSOD1) contributes to virulence on French bean, whereas a glucose oxidase (BCGOD1) is dispensable. Molecular Plant Pathology 5: 17–27.

Rossi FR, Gárriz A, Marina M, Romero M, Gonzalez, M, Collado I, Pieckenstain F. 2011. The sesquiterpene botrydial produced by *Botrytis cinerea* induces the hypersensitive response on plant tissues and its action is modulated by salicylic acid and jasmonic acid signaling. Molecular Plant-Microbe Interactions 24: 888–896.

Ryder LS, Talbot NJ. 2015. Regulation of appressorium development in pathogenic fungi. Current Opinion in Plant Biology 26: 8–13.

Sapmak A, Boyce KJ, Andrianopoulos A, Vanittanakom N. 2015. The pbrB gene encodes a laccase required for DHN-Melanin synthesis in conidia of *Talaromyces Penicillium marneffei*. PLoS One 104: e0122728.

Schouten A, Van Baarlen P, van Kan JA. 2008. Phytotoxic Nep1‐like proteins from the necrotrophic fungus *Botrytis cinerea* associate with membranes and the nucleus of plant cells. New Phytologist 177: 493–505.

Schumacher J. 2016. DHN melanin biosynthesis in the plant pathogenic fungus *Botrytis cinerea* is based on two developmentally regulated key enzyme (PKS)‐encoding genes. Molecular Microbiology 99: 729–748.

Segmüller N, Kokkelink L, Giesbert S, Odinius D, van Kan JA, Tudzynski P. 2008. NADPH oxidases are involved in differentiation and pathogenicity in *Botrytis cinerea*. Molecular Plant-Microbe Interactions 21: 808–819.

Seifert GJ. 2018. Fascinating fasciclins: a surprisingly widespread family of proteins that mediate interactions between the cell exterior and the cell surface. International Journal of Molecular Sciences 19: 1628.

Shlezinger N, Minz A, Gur Y, Hatam I, Dagdas Y, Talbot N, Sharon A. 2011. Antiapoptotic machinery protects the necrotrophic fungus *Botrytis cinerea* from host-induced apoptotic-like cell death during plant infection. PLoS Pathogens 7: e1002185.

Sharman S, Heale JB. 1977. Penetration of carrot roots by the grey mould fungus *Botrytis cinerea* Pers. ex Pers. Physiological Plant Pathology 10: 63–71.

Siegmund U, Viefhues A. 2016. Reactive Oxygen Species in the *Botrytis* – Host Interaction. In: Fillinger S, Elad Y, eds. Botrytis-The Fungus, the Pathogen and Its Management in Agricultural Systems. Cham, Switzerland: Springer: 269–290.

Simon A, Biot E. 2010. ANAIS: Analysis of NimbleGen Arrays Interface. Bioinformatics. 26: 2468–2469.

Simon A, Viaud M. 2018. Cross-reference table for Botrytis cinerea B0510 and T4 gene ids. https://doi.org/10.15454/IHYJCX,PortailDataINRAE,V2,UNF:6:PgcjbcKsoyoUHnNO6LX9Vg==[fileUNF]

Smith VL, Punja ZK, Jenkins SF. 1986. A histological study of infection of host tissue by *Sclerotium rolfsii*. Phytopathology 76: 755–759.

Stewart A, Backhouse D, Sutherland PW, Fullerton RA. 1989. The Development of Infection Structures of *Sclerotium cepivorum* on Onion. Journal of Phytopathology 126: 22–32.

Sun LL, Li M, Suo F, Liu XM, Shen EZ, Yang B, Dong MQ, He WZ, Du LL. 2013. Global analysis of fission yeast mating genes reveals new autophagy factors. PLoS Genetics 9: e1003715.

Tariq VN, Jeffries P. 1984. Appressorium formation by *Sclerotinia sclerotiorum*: scanning electron microscopy. Transactions of the British Mycological Society 82: 645–651.

Ten Have A, Espino JJ, Dekkers E, Van Sluyter SC, Brito N, Kay J, González C, van Kan JA. 2010. The *Botrytis cinerea* aspartic proteinase family. Fungal Genetics and Biology 47: 53–65.

Upadhya R, Baker LG, Lam WC, Specht CA, Donlin MJ, Lodge JK. 2018. *Cryptococcus neoformans* Cda1 and Its Chitin Deacetylase Activity Are Required for Fungal Pathogenesis. mBio 9: e02087–18.

Van den Brink J, de Vries RP. 2011. Fungal enzyme sets for plant polysaccharide degradation. Applied Microbiology and Biotechnology 91: 1477.

Van den Heuvel J, Waterreus LP. 1983. Conidial concentration as an important factor determining the type of prepenetration structures formed by *Botrytis cinerea* on leaves of French bean (*Phaseolus vulgaris*). Plant Pathology 32: 263–272.

Van Kan JA, Stassen JHM, Mosbach A, Van der Lee TAJ, Faino L, Farmer AD, Papasotiriou DG, Zhou SG, Seidl MF, Cottam E et al. 2017. A gapless genome sequence of the fungus *Botrytis cinerea*. Molecular Plant pathology 18: 75–89.

Viefhues A, Heller J, Temme N, Tudzynski P. 2014. Redox Systems in *Botrytis cinerea*: Impact on Development and Virulence. Molecular Plant-Microbe Interactions 27: 858–874.

Viefhues A, Schlathoelter I, Simon A, Viaud M, Tudzynski P. 2015. Unraveling the function of the response regulator BcSkn7 in the stress signaling network of *Botrytis cinerea*. Eukaryotic Cell 14: 636–651.

Xie M, Zhao X, Lü Y, Jin C. 2020. Chitin deacetylases Cod4 and Cod7 are involved in polar growth of *Aspergillus fumigatus*. MicrobiologyOpen 11: e943.

Yu J-H, Hamari Z, Han K-H, Seo J-A, Reyes-Domínguez Y, Scazzocchio C. 2004. Double-joint PCR: a PCR-based molecular tool for gene manipulations in filamentous fungi. Fungal Genetics and Biology 41: 973–981.

Zhang L, Wu MD, Li GQ, Jiang DH, Huang HC. 2010. Effect of mitovirus infection on formation of infection cushions and production of some virulence factors by *Botrytis cinerea*. Physiology and Molecular Plant Pathology 75: 71–80.

Zhang Y, Zhang Y, Qiu D, Zeng H, Guo L, Yang X. 2015. BcGs1, a glycoprotein from *Botrytis cinerea*, elicits defence response and improves disease resistance in host plants. Biochemical and biophysical research communications 457: 627–634.

Zhang W, Corwin JA, Copeland DH, Feusier J, Eshbaugh R, Cook DE, Atwell S, Kliebenstein DJ. 2019. Plant-necrotroph co-transcriptome networks illuminate a metabolic battlefield. eLife 8: e44279.

Zhu W, Ronen M, Gur Y, Minz-Dub A, Masrati G, Ben-Tal N, Sharon I, Savidor A, Eizner E, Valerius O et al. 2017. BcXYG1, a secreted xyloglucanase from *Botrytis cinerea* induces cell death and triggers plant defense. Plant Physiology 175: 438–456.

