## Supplemental Figures for "The infection cushion: a fungal “weapon” of plant-biomass destruction": Supplemental_Files.pdf

bioRxiv
THE PREPRINT SERVER FOR BIOLOGY

**Supporting Information**

Article title: **The infection cushion: a fungal "weapon" of plant-biomass destruction**

Authors: Mathias Choquer, Christine Rascle, Isabelle R Gonçalves, Amélie de Vallée, Cécile Ribot, Elise Loisel, Pavlé Smilevski, Jordan Ferria, Mahamadi Savadogo, Eytham Souibgui, Marie-Josèphe Gagey, Jean-William Dupuy, Jeffrey A Rollins, Riccardo Marcato, Camille Noûs, Christophe Bruel and Nathalie Poussereau

The following Supporting Information is available for this article:

**Fig. S1 *In vitro* production of infection cushions of *Botrytis cinerea.***

**Fig. S2 Gene ontology (GO) term enrichment analysis of genes differentially expressed in the infection cushion of *Botrytis cinerea*.**

**Fig. S3 Regulation of secondary metabolism biosynthesis key genes in the infection cushion of *Botrytis cinerea.***

**Fig. S4 Putative secondary metabolism gene clusters up-regulated in the infection cushion of *Botrytis cinerea*.**

**Fig. S5 Mutagenesis of genes encoding fasciclin-like proteins in *Botrytis cinerea.***

**Fig. S6 Comparison of the *Botrytis cinerea* infection cushion (IC) transcriptome and the transcriptomes of four IC-deficient mutants.**

**Table S1 Results of the microarray analysis of the infection cushion of *Botrytis cinerea*.**

**Table S2 RT-qPCR validation of microarray expression profiles from the infection cushion of *Botrytis cinerea* produced *in vitro.***

**Table S3 Up-accumulated proteins in the secretome of the infection cushion of *Botrytis cinerea.***

**Table S4 Constructs and primers used in this study.**

bioRχiv
THE PREPRINT SERVER FOR BIOLOGY

**Fig. S1 *In vitro* production of infection cushions of *Botrytis cinerea*.** (a) Light microscopy of mycelia produced from conidia of *B. cinerea* spread onto PDA plates overlaid with cellophane and culture for 44h. (b) Light microscopy of mycelia produced from conidia inoculated in PDB-containing flasks and cultured under agitation for 44h. Magnifications are indicated.

**Supplementary Figure S1**

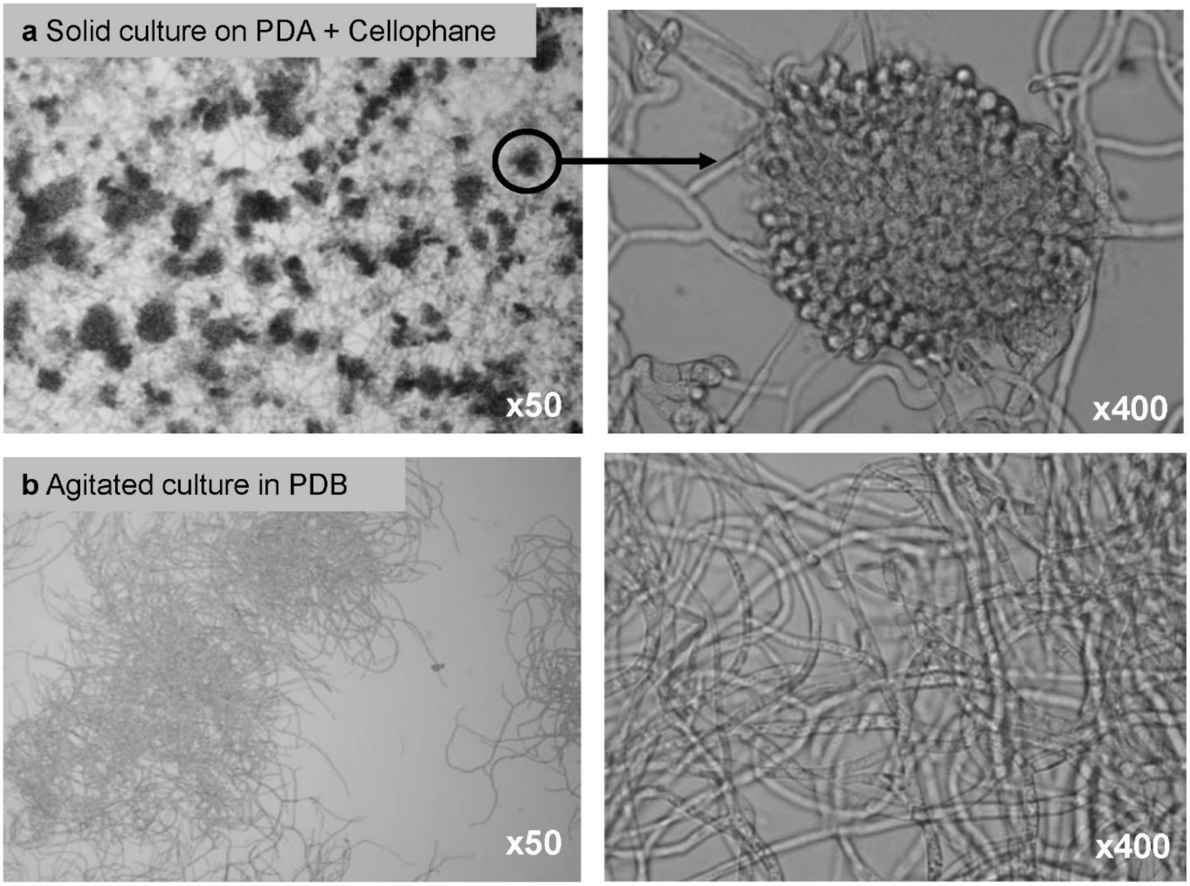

bioRxiv
THE PREPRINT SERVER FOR BIOLOGY

**Fig. S2 Gene Ontology (GO) term enrichment analysis of genes differentially expressed in the infection cushion of *Botrytis cinerea*.** Significantly (Fisher's exact test; p-values < 0.05) over-represented biological processes (BP) for the (a) 1,231 genes up-regulated in IC and (b) 1,422 genes down-regulated in IC. Enriched categories are classified according to p-values. Ratios inside the pie charts indicate the number of differentially expressed genes (underlined) relative to the total number of genes in each BP category (*italics*).

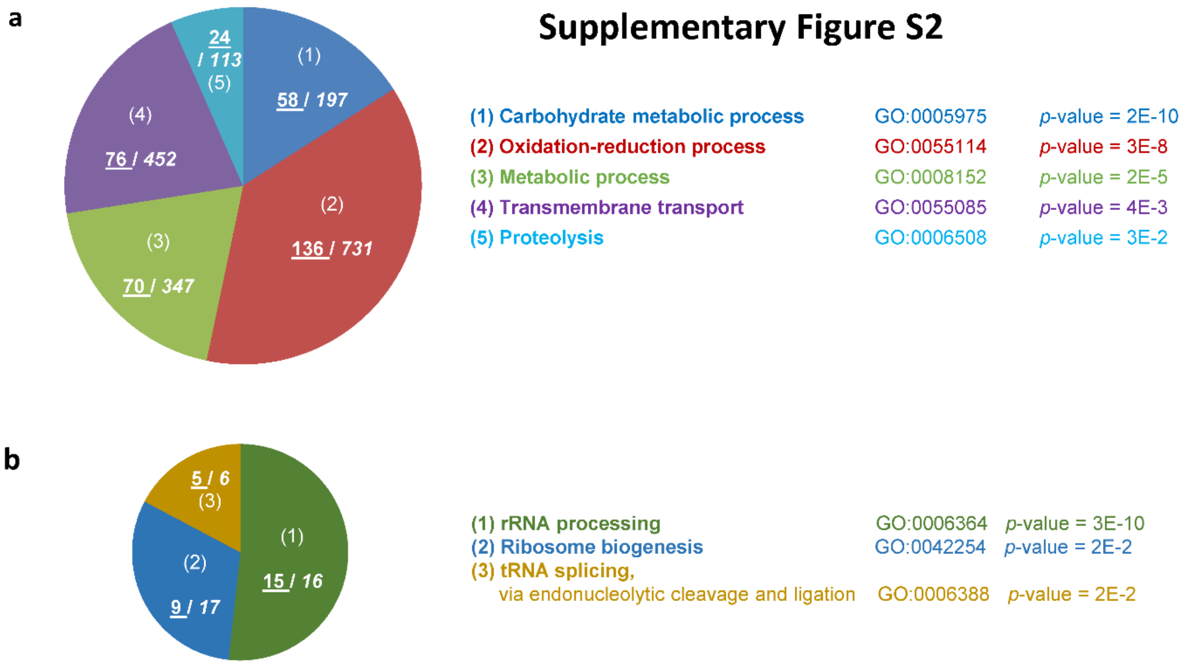

bioRxiv
THE PREPRINT SERVER FOR BIOLOGY

**Fig. S3 Regulation of secondary metabolism biosynthesis key genes in the infection cushion of *Botrytis cinerea*.** (a) Inventory of secondary metabolism (SM) biosynthesis key enzymes. The names, acronyms and number of genes coding for the SM key enzymes in *B. cinerea* are presented. (b) Hierarchical clustering of the expression of the 42 predicted SM key enzymes-encoding genes in IC (IC, 4 replicates) and control mycelium (Myc, 3 replicates). The normalized expression intensities are clustered and represented by color-coded squares; Shades of green and red depict down- and up-regulation in IC, respectively (Fold change ≤ -2 or ≥ 2 and FDR<0.05). Background corresponds to genes considered not expressed (with normalized intensities lower than the 99th percentile of random probes hybridization signals in all biological replicates).

bioRxiv
THE PREPRINT SERVER FOR BIOLOGY

### Supplementary Figure S3

**a**

| Key enzymes | Polyketide synthases | Non-ribosomal peptide synthetases | PKS-NRPS hybrids | Sesquiterpene cyclases | Diterpen cyclases | Dimethylallyl tryptophan synthases | Other |
|---|---|---|---|---|---|---|---|
| Acronym | PKS1, PKS2, PKS8 to PKS21 | NRPS1 to NRPS9 | PKS3 to PKS7 | STC1 to STC5 | DTC1 & DTC3 | DMATS, DMATS2 | CHS1, PAX1, PHS1 |
| Number of genes | 16 | 9 | 5 | 5 | 2 | 2 | 3 |

**b**

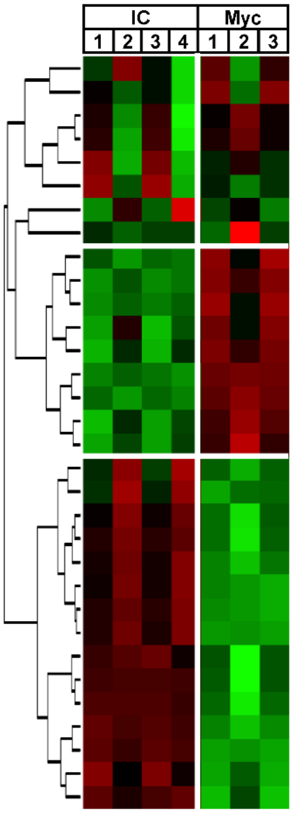


| Gene ID | Chip reference | FC | FDR | Name | Description |
|---|---|---|---|---|---|
| Bcin05g06220 | BofuT4_P135120 | -1.0 | $8.2^{E-1}$ | *Bcpks15* | PKS |
| Bcin01g04920 | BofuT4_P044420 | -2.2 | $3.1^{E-1}$ | *Bcdtc1* | DTC |
| Bcin04g03550 | BofuT4_P096230 | -1.4 | $3.4^{E-1}$ | *Bcstc4* | STC |
| Bcin16g03570 | BofuT4_P150070 | -1.7 | $3.4^{E-1}$ | *Bcnrps3* | NRPS |
| Bcin16g01940 | BofuT4_P033980 | 2.5 | $9.5^{E-1}$ | *Bcdmats1* | DMATS |
| Bcin01g11550 | BofuT4_P085990 | 1.4 | $5.8^{E-1}$ | *Bcpks5* | PKS-NRPS |
| Bcin01g04560 | BofuT4_P163490 | 1.2 | $6.4^{E-1}$ | *Bcphs1* | Phytoene synthase |
| Bcin02g02380 | BofuT4_P109840 | -1.3 | $4.7^{E-1}$ | *Bcnrps4* | NRPS |
| Bcin02g08830 | BofuT4_P088290 | -2.7 | $2.6^{E-2}$ | *Bcpks18* | PKS |
| Bcin14g01290 | BofuT4_P059830 | -10 | $4.1^{E-3}$ | *Bcpks11* | PKS |
| Bcin14g01300 | BofuT4_P059820 | -5.7 | $2.8^{E-2}$ | *Bcnrps9* | NRPS |
| Bcin12g00690 | BofuT4_P120030 | -1.6 | $1.2^{E-1}$ | *Bcnrps2* | NRPS |
| Bcin05g08400 | BofuT4_P103050 | -4.5 | $3.0^{E-2}$ | *Bcpks21* | PKS |
| Bcin01g11450 | BC1G_15493 | -24 | $7.0^{E-4}$ | *Bcnrps7* | NRPS |
| Bcin02g08770 | BofuT4_P088370 | -4.2 | $1.4^{E-3}$ | *Bcpks12* | PKS |
| Bcin08g02350 | BofuT4_P130680 | -2.5 | $2.3^{E-2}$ | *Bcstc2* | STC |
| Bcin03g04360 | BofuT4_P020280 | -3.4 | $2.9^{E-2}$ | *Bcpks3* | PKS-NRPS |
| Bcin04g00640 | BofuT4_P085340 | 3.2 | $1.3^{E-1}$ | *Bcpks20* | PKS |
| Bcin01g03520 | BofuT4_P162160 | 7.0 | $9.6^{E-2}$ | *Bcstc5* | STC |
| Bcin04g01390 | BofuT4_P115630 | 2.8 | $3.3^{E-2}$ | *Bcnrps5* | NRPS |
| Bcin10g00040 | BofuT4_P071110 | 8.5 | $1.7^{E-2}$ | *Bcpks7* | PKS-NRPS |
| Bcin08g00290 | BofuT4_P119530 | 13 | $1.2^{E-2}$ | *Bcpks19* | PKS |
| Bcin11g02650 | BofuT4_P037730 | 96 | $1.1^{E-2}$ | *Bcnrps8* | NRPS |
| Bcin08g03560 | BofuT4_P132450 | 13 | $3.7^{E-3}$ | *Bcdtc3* | DTC |
| Bcin11g02700 | BofuT4_P037790 | 50 | $5.2^{E-3}$ | *Bcpks4* | PKS-NRPS |
| Bcin13g02130 | BofuT4_P125060 | 1.5 | $3.8^{E-2}$ | *Bcchs1* | Chalcone synthase |
| Bcin12g06390 | BCexp3_CND15 | 4.2 | $2.7^{E-2}$ | *Bcbot2* | STC1 |
| Bcin01g00090 | BC1G_15837 | 6.6 | $1.1^{E-2}$ | *Bcboa9* | PKS9 |
| Bcin01g00060 | BC1G_16087 | 12 | $1.6^{E-3}$ | *Bcboa6* | PKS-NRPS(PKS6) |
| Bcin07g02920 | BofuT4_P119030 | 10 | $5.9^{E-4}$ | *Bcpks8* | PKS |
| Bcin03g08050 | BofuT4_P077210 | 5 | $2.4^{E-2}$ | *Bcpks13* | PKS |
| Bcin01g03730 | BofuT4_P162470 | 1.9 | $1.3^{E-2}$ | *Bcnrps6* | NRPS |
| Bcin13g05830 | BofuT4_P062370 | Background | | *Bcstc3* | STC |
| Bcin14g04900 | BofuT4_P098370 | | | *Bcdmats2* | DMATS |
| Bcin14g00600 | BofuT4_P088990 | | | *Bcpks1* | PKS |
| Bcin02g01680 | BofuT4_P108860 | | | *Bcpks2* | PKS |
| Bcin13g01510 | BofuT4_P125860 | | | *Bcpks10* | PKS |
| Bcin16g01830 | BofuT4_P034180 | | | *Bcpks14* | PKS |
| Bcin16g05040 | BofuT4_P152140 | | | *Bcpks16* | PKS |
| Bcin03g02010 | BofuT4_P052990 | | | *Bcpks17* | PKS |
| Bcin12g04980 | BofuT4_P002220 | | | *Bcnrps1* | NRPS |
| Bcin05g05670 | BofuT4_P135790 | | | *BcPax1* | PaxC-like terpen synthase |

bioRχiv
THE PREPRINT SERVER FOR BIOLOGY

**Fig. S4 Putative secondary metabolism gene clusters up-regulated in the infection cushion of *Botrytis cinerea.*** (a) Hierarchical clustering of the expression of SM key enzymes-encoding genes and surrounding genes in IC (IC, 4 replicates) and control mycelium (Myc, 3 replicates). The normalized expression intensities are represented by color-coded squares; Shades of green and red depict down- and up-regulation in IC, respectively (Fold change ≤ -2 or ≥ 2 and FDR<0.05). (b) Genomic localization and representation of the 4 putative SM gene clusters DTC3, PKS7, PKS8 and PKS4-NRPS8. The genes up-regulated in IC are indicated in red and the supposed ends of each cluster are represented by the first neighboring genes non-regulated in IC (grey).

bioRxiv
THE PREPRINT SERVER FOR BIOLOGY

### Supplementary Figure S4

**a**

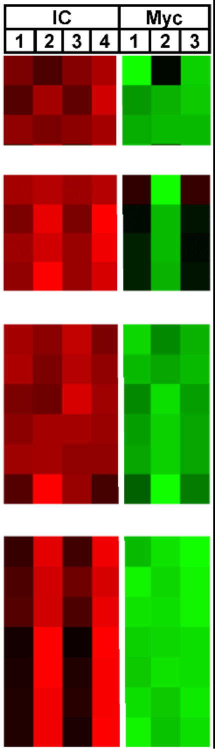


| Gene ID | Chip reference | FC | FDR | Name | Description |
|---|---|---|---|---|---|
| **Dtc3 cluster** | | | | | |
| Bcin08g03550 | BofuT4_P132440 | 3.4 | 2.9E-2 | no | Hypothetical protein |
| Bcin08g03560 | BofuT4_P132450 | 12 | 3.7E-3 | *Bcdtc3* | **Diterpen cyclase** |
| Bcin08g03570 | BofuT4_P132480 | 5.1 | 2.1E-4 | no | **Cytochrome P450** |
| **Pks7 cluster** | | | | | |
| Bcin10g00030 | BofuT4_P071050 | 2.9 | 1.1E-1 | no | hypothetical protein |
| Bcin10g00010 | BC1G_16236 | 16 | 3.1E-2 | no | **hypothetical protein** |
| Bcin10g00020 | BC1G_16237 | 19 | 1,6E-2 | no | **MFS transporter** |
| Bcin10g00040 | BofuT4_P071110 | 8.5 | 1.6E-2 | *Bcpks7* | **PKS-NRPS hybrid** |
| **Pks8 cluster** | | | | | |
| Bcin07g02900 | BofuT4_P118990 | 9.6 | 1.5E-3 | no | **ABC transporter** |
| Bcin07g02920 | BofuT4_P119030 | 10 | 5.9E-4 | *Bcpks8* | **Polyketide synthase** |
| Bcin07g02930 | BofuT4_P119040 | 2.9 | 2.9E-3 | no | Hypothetical protein |
| Bcin07g02910 | BofuT4_P119010 | 7.0 | 5.6E-4 | no | EF-hand protein |
| Bcin07g02890 | BofuT4_P118980 | 8.3 | 4.1E-4 | no | Hypothetical protein |
| Bcin07g02940 | BofuT4_P119050 | 2.0 | 2.4E-2 | no | Kelch repeat-containing protein |
| **Nrps8/Pks4 cluster** | | | | | |
| Bcin11g02650 | BofuT4_P037730 | 95 | 1.1E-2 | *Bcnrps8* | **Non-ribosomal peptide synthetase** |
| Bcin11g02670 | BofuT4_P037750 | 27 | 3.5E-3 | no | **Dehydrogenase/reductase** |
| Bcin11g02700 | BofuT4_P037790 | 50 | 5.2E-3 | *Bcpks4* | **Polyketide synthase** |
| Bcin11g02690 | BofuT4_P037770 | 51 | 2.7E-2 | no | EF-hand protein |
| Bcin11g02640 | BofuT4_P037720 | 56 | 2.0E-2 | no | **Monooxygenase** |
| Bcin11g02680 | BofuT4_P037760 | 131 | 1.6E-2 | no | Acetyl-coA synthetase |
| Bcin11g02660 | BofuT4_P037740 | 32 | 2.1E-2 | no | **Methyl transferase** |

**b**

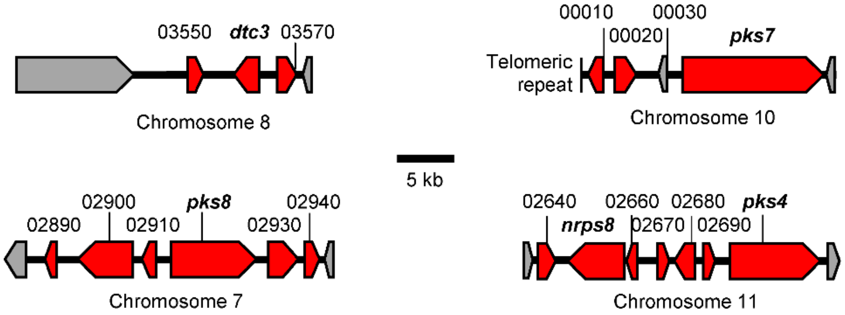

bioRxiv
THE PREPRINT SERVER FOR BIOLOGY

**Fig. S5 Mutagenesis of genes encoding fasciclin-like proteins in *Botrytis cinerea.*** (a) Creation of the Δ*Bcflp1* deletion mutant (left) and Southern analysis (right). A split-marker strategy was used to replace the *Bcflp1* gene of the *B. cinerea* B05.10 strain by a hygromycin resistance gene (Hygro). The promoter and terminator DNA regions of *Bcflp1* (5'- and 3'-Flank) were inserted to the overlapping semi-replacement DNA cassettes in order to target them to the appropriate genomic locus. Homologous recombination (dashed lines) led to gene replacement. Genomic DNA of two hygromycin resistant clones (T2, T3) and the parental strain (WT) was digested by EcoRI (E) and subjected to Southern blot analysis using the 3'-flanking region as DNA probe (blue bar). (b) Creation of the Δ*Bcflp2* deletion mutant (left) and Southern analysis (right). Following the same strategy described in (a), six transformants (T6-12) were analyzed by using the 5'-flanking region as DNA probe (green bar). (c) Creation of the Δ*Bcflp1::*Δ*Bcflp2* double deletion mutant (left) and Southern analysis (right). The same strategy described in (a) was used with the nourseothricin resistance gene (Nourseo) replacing the hygromycin resistance gene and the Δ*Bcflp1* strain being the recipient strain for the transformation experiment. Three transformants (T) were analyzed by using the 3'-flanking region as DNA probe (blue bar). Gene replacements in the deletion strains were also verified by diagnostic PCR (data not shown).

bioRχiv
THE PREPRINT SERVER FOR BIOLOGY

### Supplementary Figure S5

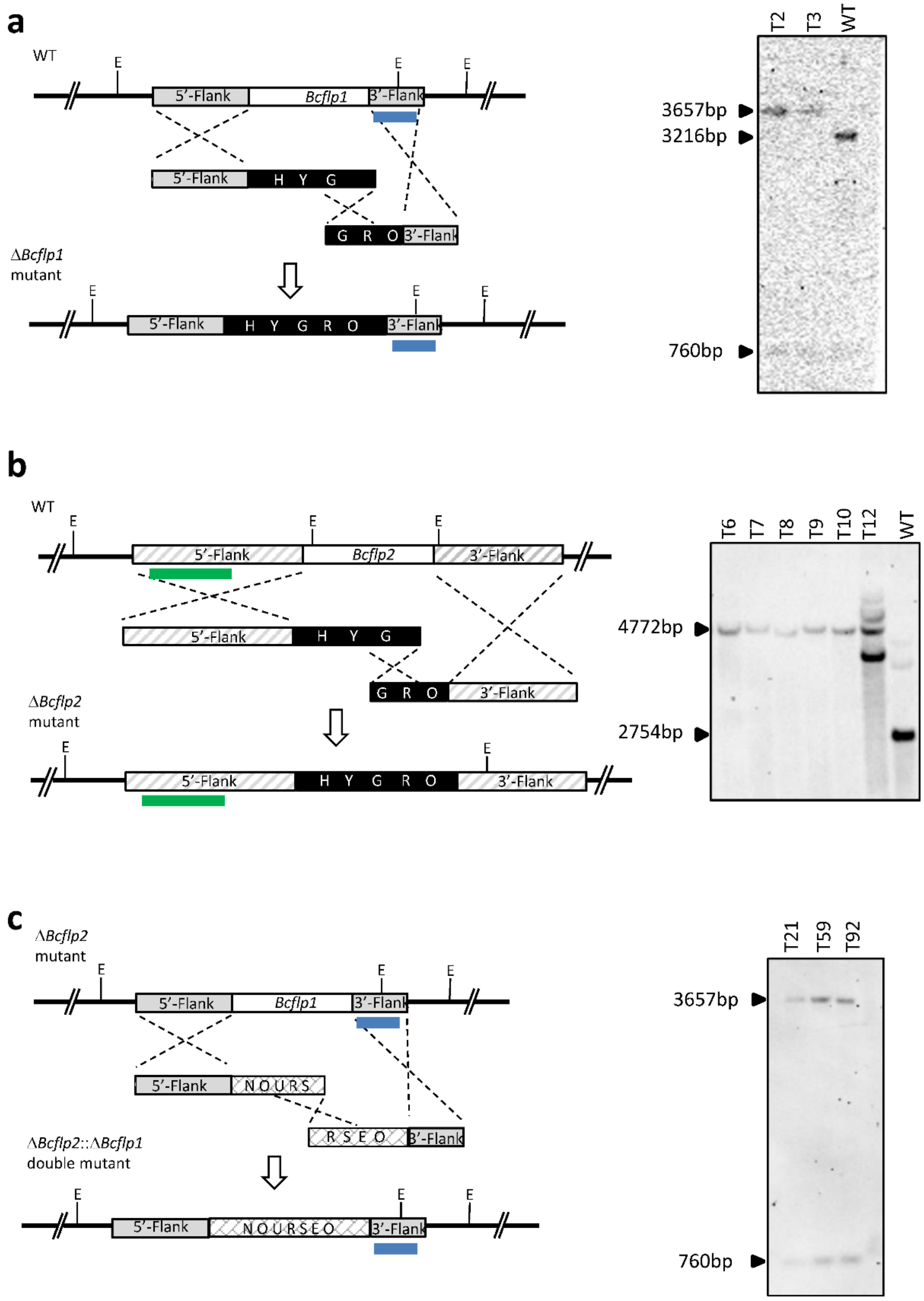

bioRxiv
THE PREPRINT SERVER FOR BIOLOGY

**Fig. S6 Comparison of the *Botrytis cinerea* infection cushion (IC) transcriptome and the transcriptomes of four IC-deficient mutants.** The transcriptomes (RNAseq) of four *B. cinerea* mutants impaired in the formation of IC were published (De Vallée *et al.*, 2019) and used for data comparison with the transcriptome of IC (this study). Genes repressed in these IC-deficient mutants were assumed to be induced in the infection cushion of the wild-type strain. This inverted correlation was confirmed for 40 genes among the top 100 upregulated genes in IC as they showed a common down-regulation in the four IC-deficient mutants.

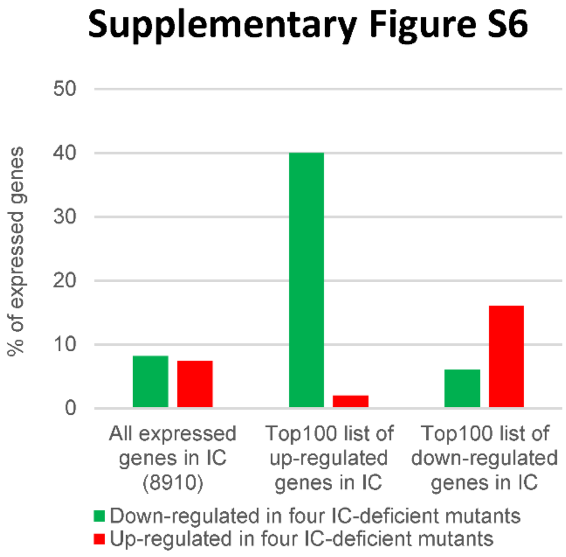

bioRxiv
THE PREPRINT SERVER FOR BIOLOGY

**Table S1 Results of the microarray analysis of the infection cushion of *Botrytis cinerea*.** Table showing all Bcin genes analyzed by the Nimblegen chip (n=11134), including Bcin genes up-regulated in IC (n=1231), Bcin genes down-regulated in IC (n=1422), Bcin genes expressed but not differentially regulated in IC (n=6757), and Bcin genes not expressed in IC nor mycelium control (n=1724). Bcin genes correspond to the genome of B05.10 strain structurally annotated by Van Kan *et al.* (2017) and available at http://fungi.ensembl.org/Botrytis_cinerea. For each Bcin gene, the indicated chip reference is the corresponding gene from two previous genome annotations BofuT4 or BC1G (Amselem *et al*., 2011). Usual names were imported from the *Botrytis cinerea* Portal (http://botbioger.versailles.inra.fr/botportal/). Fold change is the ratio between the mean of IC normalized intensities and the mean of mycelium control normalized intensities. ANOVA p-values were submitted to a False Discovery Rate correction (FDR). Normalized intensities are given for the infection cushion (4 biological replicates) and the mycelium control (3 biological replicates). Presence for a putative signal peptide was predicted using the SignalP 5.0 Server (Almagro Armenteros *et al*., 2019). Transcripts with a corrected p-value < 0.05 and for which a fold change ≤ -2 or ≥ 2 was observed between the two conditions were considered to display significant differential expression. Genes were considered not expressed when their normalized intensity was smaller than the 99th percentile of random probes hybridization signals in all biological replicates. For IC replicates 1 to 4, the 99th percentile of random probes hybridization signals was: 752, 572, 532 and 995, respectively. For mycelium replicates 1 to 3, the 99th percentile of random probes hybridization signals was: 752, 634 and 599, respectively (as determined by R software).

bioRxiv
THE PREPRINT SERVER FOR BIOLOGY

**Table S2** ***In vitro*** **RT-qPCR validation of microarray expression profiles from the infection cushion of *Botrytis cinerea*.** Forty genes were chosen to test the differential expression recorded by the microarray analysis in the IC versus control mycelium. New biological replicates (n=3) were prepared and the expression of 24 up-regulated genes, 6 down-regulated genes and 10 non-regulated genes was measured by RT-qPCR. Gene expression levels were calculated following the $2^{-\Delta\Delta CT}$ method using constitutively expressed elongation factor *Bcef1α* as a reference (the use of the house-keeping actin (*BcactA*) and pyruvate dehydrogenase (*Bcpda1*) genes gave similar results; data not shown). Genes are ordered in the table according to their fold change in the microarray analysis and color codes corresponding to different fold change intensities are indicated. FC, fold change; FDR, false discovery rate; DE, differential expression.

bioRχiv
THE PREPRINT SERVER FOR BIOLOGY

**Table S3 Up-accumulated proteins in the secretome of the infection cushion of *Botrytis cinerea.*** Comparison of the secretome of IC versus vegetative mycelium revealed 79 proteins up-accumulated. For quantification, all unique peptides of an identified protein were included, and the total cumulative abundance was calculated by summing the abundances of all peptides allocated to the respective protein. ANOVA test was applied at the protein level. Three independent biological experiments were conducted and analyzed. The mass spectrometry proteomics data have been deposited to the ProteomeXchange Consortium via the PRIDE partner repository with the dataset identifier PXD016885. Gene ID and protein accession numbers can be found in Ensembl Fungi release database (http://fungi.ensembl.org/Botrytis_cinerea/Info/Index). The ratios between the average protein abundances in the IC-enriched and vegetative mycelia are indicated with the associated p-values. The up-regulated proteins, identified in 3 biological replicates with a minimum of 2 unique peptides, are listed and classified according to their functional category (manual annotation). Six known necrosis inducers are highlighted in yellow. SignalP and ApoplastP prediction softwares were respectively used to predict the presence of a signal peptide in these proteins sequence and the presence of these proteins in a plant apoplast (http://apoplastp.csiro.au/; Sperschneider *et al*., 2017). The up-accumulation is colored in red and the fold change values (FC) are indicated with the associated p-values. The microarray data (Chip reference, fold change and FDR) of these proteins-encoding genes are indicated. The behaviour of these proteins and corresponding genes in 4 IC-deficient mutants (De Vallée *et al*., 2019) are also indicated. PCWDE: Plant cell wall degrading enzymes. **Sperschneider J, Dodds PN, Singh KB, Taylor JM. 2018.** APOPLASTP: prediction of effectors and plant proteins in the apoplast using machine learning. *New Phytologist*. **217:** 1764-1778.

bioRχiv
THE PREPRINT SERVER FOR BIOLOGY

**Table S4 Primers used in this study for RT-qPCR analysis and constructs.** For *Bcflp1* deletion construct, the 5'- and 3'- flanking regions of *Bcflp1* (1.115 kb and 0.624 kb) were amplified from *B. cinerea* genomic DNA using the primers pairs flp1-H1/flp1-H2 and flp1-H3/flp1-H4, respectively. flp1-H2 and flp1-H3 also contained sequences homologous to the hygromycin resistance cassette containing the hph gene under control of the oliC promoter. This cassette was generated using primers Hyg-H5 and Hyg-H6 and plasmid pFV8 as a template (Villalba *et al*., 2008). The purified amplicons were fused in a second round of PCR without any primer. Finally, a third step of PCR led to the amplification of two overlapping semi- replacement cassettes using nested primers flp1-H7/HYG-H7 and GRO-H8/flp1-H8 respectively. For *Bcflp2* deletion construct, the same strategy was applied using primers pairs flp2-H1/flp2-H2 for 5'- flanking region of *Bcflp2* (1.962 kb) and flp2-H3/flp2-H4 for 3'- flanking region of *Bcflp2* (1.468 kb) and resulted in two overlapping semi- replacement cassettes using nested primers flp2-H7/HYG-H7 and GRO-H8/flp2-H8 respectively. The development of double replacement mutants was achieved by transforming the previously generated *ΔBcflp2* mutant. Double deletion mutants *ΔBcflp2::ΔBcflp1* were constructed with the same strategy excepted that nourseothricin resistance was used instead of hygromycin. The 5'- and 3'- flanking regions of *Bcflp1* (1.095 kb and 0.56 kb) were amplified from *B. cinerea* genomic DNA using the primers pairs flp1-N1/flp1-N2 and flp1-N3/flp12-N4, respectively. The nourseothricin resistance cassette was amplified using flp1-N5 and flp1-N6 primers before fusion with the 5' and 3' flanking regions. Finally, two semi-cassettes were amplified using nested primers pairs flp1-N7/NOU-N7 and URS-N8/flp1-N8 respectively. *Bcflp1* replacement by the hygromycin cassette was verified using the primers pairs flp1-F5'/ hygro5' and hygro3'/flp1-F3' and the primers pairs flp1-N4/nour5' and nour3'/flp1-N1 for *Bcflp1* replacement by the nourseothricin cassette. *Bcflp2* replacement by the hygromycin cassette was verified using the primers pairs flp2-F5'/ hygro5' and hygro3'/flp2-F3'. **Villalba F, Collemare J, Landraud P, Lambou K, Brozek V, Cirer B, Morin D, Bruel C, Beffa R, Lebrun MH. 2008.** Improved gene targeting in *Magnaporthe grisea* by inactivation of *MgKU80* required for non-homologous end joining. *Fungal Genetics and Biology*. **45:** 68-75.
